# Monte-Carlo Diffusion-Enhanced Photon Inference: Distance Distributions And Conformational Dynamics In Single-Molecule FRET

**DOI:** 10.1101/385252

**Authors:** Antonino Ingargiola, Shimon Weiss, Eitan Lerner

**Affiliations:** Department of Chemistry and Biochemistry, University of California Los Angeles, USA; Department of Biological Chemistry, The Alexander Silberman Institute of Life Sciences, The Hebrew University, Jerusalem, Israel

## Abstract

Single-molecule Förster Resonance Energy Transfer (smFRET) is utilized to study the structure and dynamics of many bio-molecules, such as proteins, DNA and their various complexes. The structural assessment is based on the well-known Förster relationship between the measured efficiency of energy transfer between a donor (D) and an acceptor (A) dye and the distance between them. Classical smFRET analysis methods called photon distribution analysis (PDA) take into account photon shot-noise, D-A distance distribution and, more recently, interconversion between states in order to extract accurate distance information. It is known that rapid D-A distance fluctuations on the order of the D lifetime (or shorter) can increase the measured mean FRET efficiency and thus decrease the estimated D-A distance. Nonetheless, this effect has been so far neglected in smFRET experiments, potentially leading to biases in estimated distances.

Here we introduce a PDA approach dubbed Monte-Carlo-diffusion-enhanced photon inference (MC-DEPI). MC-DEPI recolor detected photons of smFRET experiments taking into account dynamics of D-A distance fluctuations, multiple interconverting states and photo-blinking. Using this approach, we show how different underlying conditions may yield identical FRET histograms and how the additional information from fluorescence decays helps distinguishing between the different conditions. We also introduce a machine learning fitting approach for retrieving the D-A distance distribution, decoupled from the above-mentioned effects. We show that distance interpretation of smFRET experiments of even the simplest dsDNA is nontrivial and requires decoupling the effects of rapid D-A distance fluctuations on FRET in order to avoid systematic biases in the estimation of the D-A distance distribution.

## 1 Introduction

Förster resonance energy transfer (FRET) is a phenomenon in which an electronically excited fluorophore transfers a fraction of its excitation energy non-radiatively to another fluorophore in the ground-state. The energy transfer occurs as long as the fluorescence spectrum of the first fluorophore, the donor (D), overlaps with the excitation spectrum of the second fluorophore, the acceptor (A), and as long as they are in close proximity. The efficiency of FRET depends on the sixth power of the distance between the D and A fluorophores.^1^ This distance dependence gained FRET the term “molecular ruler” and indeed it has been used as such to resolve conformational information for inter-fluorophore distances in the range 2-10 nm on biological molecules since 1967.^2^ Yet, such measurements at the ensemble level allowed retrieval of ensemble-averaged distances that may have been interpreted in many different ways.

The capability to measure FRET at the single molecule level allowed classification of molecules of the ensemble into classes with different FRET efficiencies, as well as identifying the dynamics of interconversion between them. Indeed, since single-molecule FRET (smFRET) was introduced in 1996,^3^ it has been used by many different labs to report conformational sub-populations and dynamics of a myriad of biological macromolecules, such as RNA, DNA, proteins and their complexes.^4^ Each of these sub-populations of FRET efficiencies represents a time-average over a few milliseconds of either a single conformational state or of multiple different conformational states with transitions between them that occur faster than this timescale. Therefore each FRET sub-population entails the distance information on a conformational state or on a millisecond average of multiple rapidly interconverting conformational states.^4–10^

In the last few years smFRET has gradually become a tool used for retrieval of intramolecular distance information.^4^ Performing such measurements on a biological molecule doubly labeled at many different pairs of atomic positions has opened the door for determination of the ensemble of structures that define different conformational states existing in the solution at ambient temperature.^5,11–14^ However, there can still be many different distance-related interpretations of a FRET sub-population, hence the retrieval of distance information from smFRET sub-populations is far from being straightforward.

For the task of standardizing the procedure of distance information retrieval from sm-FRET measurements, FRET between dyes attached to a simple rigid molecule such as ds-DNA has been used.^15,16^ However, even in such a simple molecule, the distance interpretation of an smFRET measurement may be complicated by: (1) the distribution of the D-A distance,^17,18^ (2) how fast does this distance change^19,20^ and (3) how do photophysical processes that compete with FRET influence the outcome of the smFRET measurements.^21–23^

A single conformational state can be described by a potential well depending on the D-A distance as a reaction coordinate^24^ and on a D-A self-diffusion coefficient, which describes the distance dynamics in the potential well.^24,25^ Therefore, a conformational state can be defined by the distance distribution at equilibrium *p_Eq._*(*r*), and the inter-dye self-diffusion coefficient *D*. It is also known that distance changes occurring in the timescale of the D fluorescence decay or faster, lead to enhancement of the FRET efficiency relative the static case.^20,24–27^ Additionally, different conformational states interconverting on time scales of 10 μs (or faster), yield a single shot-noise limited millisecond-averaged FRET peak. Finally, there are three well-known sources of bias in smFRET measurements: (1) the *γ*-factor, accounting for the imbalance between the detected D and A signal due to differences fluorescence quantum yields and detection efficiencies, (2) the donor spectral leakage into the acceptor detection channel and (3) acceptor photons due to direct excitation by the donor laser.^28^ For all these reasons, different combinations of distance distribution and diffusion coefficient may yield the same millisecond-averaged FRET sub-population.

Additional experimental information is beneficial to circumvent these difficulties and to more accurately retrieve distance information. These may include fluorescence anisotropy decays and their usage in smFRET when using pulsed excitation and time-correlated single photon counting (TCSPC),^29–31^ fluorescence correlation spectroscopy (FCS)^32–34^ and other methods that rely on photon statistics such as probability distribution analysis (PDA),^6,35^ burst variance analysis (BVA)^5,7,8^ or two-channel kernel-based density distribution estimator (2CDE).^9^ In summary, a multi-parameter approach may produce enough experimental data to retrieve the underlying distance information accurately, decoupling the results from all other possible effects.^31^

In principle, the model that will best fit the experimental results should yield modeled photons with exactly the same features as the experimental ones, namely their detection time relative to the beginning of the measurement, relative to the excitation moment and the channel at which they were detected (D or A fluorescence). Therefore, the richest information source is found in the detected photons themselves, the time intervals between them and their identity (donor or acceptor photons). A photon-by-photon approach may retrieve the maximal information content.^36^ In that context, a PDA approach consists in “re-coloring” the donor and acceptor photons of single-molecule bursts according to an underlying model. Re-coloring is performed by a Monte-Carlo (MC) simulation of photon numbers in each burst, generated by a Binomial distribution. Then, histograms from the re-colored bursts are compared to the the experimental ones. This approach has been extensively used to fit experimental histograms of FRET efficiency broadened beyond shot-noise by dynamics in a single distance distribution^37^ or by dynamics of interconversion between two (or more) FRET-related conformational states (without specification of their underlying distance distributions^6,38^). However, PDA implementations have so far neglected the FRET-enhancement occurring due to picosecond-to-nanosecond D-A distance changes. Crucially, the single fixed FRET efficiency assigned to each burst in PDA, implicitly assumes a broadening of the FRET peak due to static heterogeneity, while, in most cases, it is more realistic to assume fast D-A distance fluctuations due to linker dynamics. Finally, while PDA has been used mainly for fitting experimental FRET histograms, a similar approach can be used to fit experimental results presented in other types of histograms, also derived from smFRET measurements (e.g. fluorescence decays).

In this work, we introduce MC-diffusion-enhanced photon inference (MC-DEPI), a photon-by-photon Monte-Carlo-based re-coloring approach to properly and accurately analyze sm-FRET experiments, taking into account D-A self diffusion and other effects (single or multiple interconverting states, photo-blinking, correction factors). In MC-DEPI, we model D-A self diffusion trajectories as a stochastic process with a characteristic relaxation time *τ_relax_*. Instantaneous D-A distances can be computed at arbitrary time points using an Ornstein-Uhlenbeck (OU) process and the Gillespie direct-update formula.^39^ Note that, while an OU process directly models Gaussian distributed D-A distances, other distance distribution can be evaluated using a simple transformation (refer to Eq. 15 in sub-section 3.2 on the dynamics module of MC-DEPI). In MC-DEPI, D-A distances are first computed at each photon timestamp, considered as the D excitation time. Then we simulate the D de-excitation process leading to either a D or an A photon. For this purpose, we simulate D-A distance trajectories with time steps much smaller than D and A fluorescence lifetimes. The de-exitation process depends on the simulated trajectories and FRET efficiencies. At the end, for each timestamp we obtain the photon color (either D or A) and the nanotime (time separation between dye excitation and photon emission). Thus, the simulated data yields FRET histograms as well as donor and acceptor fluorescence decays. In the final step, we use a sequential model-based optimization to find optimal parameters which fit the simulation to the experimental results.

This paper is organized as follows. We first discuss the dependence of FRET on D-A distances. Then, we introduce the theoretical framework behind MC-DEPI. Afterwards we show how a given FRET population may result from many different underlying conditions and how the additional information from other histograms derived from the data can be used to decipher which of the conditions describes best the underlying conformational state or states. Then, we show, as a proof of concept, the results of MC-DEPI-based analysis of nanosecond-alternating laser excitation (nsALEX)^29,30,40^ smFRET measurements of the distance between two dyes attached to a pair of bases in dsDNA. We illustrate how complex its distance-related interpretation may be and how MC-DEPI allows for an accurate retrieval of distance information. Finally, we discuss other possible uses of MC-DEPI in the analysis of more complex systems, whether due to complex photophysics or due to complex distance dynamics.

## 2 Experimental section

A full account for the methods used in this work can be found in the Supporting Information. The core MC-DEPI recoloring simulations are implemented in the open source depi python package (https://github.com/OpenSMFS/depi). The python notebooks used for this paper are available on GitHub (https://github.com/tritemio/mcdepi2018-paper-analysis). The experimental data files are available on Figshare (https://doi.org/10.6084/m9.figshare.6931271).

## 3 Results

In FRET experiments one measures *E*, the efficiency of transfer of excitation energy from a donor (D) to an acceptor (A) fluorophore. This efficiency depends on the D-A distance, *r*, according to the Förster relation (Eq. 1):

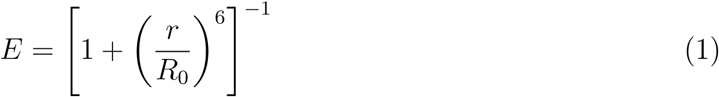

where *R*_0_, known as the Förster radius, is the distance at which the donor excitation energy is transferred to the acceptor with 50% efficiency. *R*_0_ depends on the spectral overlap between the D fluorescence spectrum, *F_D_*(*λ*), and the A extinction spectrum, *ϵ_A_*(*λ*), the D fluorescence quantum yield, *ϕ_D_*, the orientation factor of D and A fluorophores, *κ*^2^, and the refractive index of the medium between them, *n* (Eq. 2).

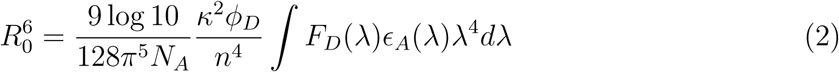

The orientation factor, *κ*^2^, is a function of the angles *θ_D_* or *θ_A_* between the direction of the D or A dipoles and the line connecting the centers of the D and A dipoles, and the angle *φ* between the D dipole and the A dipole, (Eq. 3).

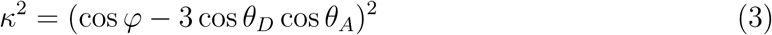

The dependence of *E* on the sixth power of *r* makes FRET a spectroscopic ruler sensitive to distances in the range between 1*/*2*R*_0_ and 3*/*2*R*_0_. Therefore, for a D-A pair with *R*_0_ of 60 Å, FRET will report accurately on distances in the range 30-90 Å.

The Jablonski diagrams depict the photophysical processes that occur following the excitation of the D fluorophore. After D is excited from the *S*_0_ ground-state to the *S*_1_ excited-state (with a rate *k_D,ex_*), it will be de-excited back to *S*_0_, *S*_1_ → *S*_0_, with a rate, *k_D_*, either radiatively yielding an emitted photon or nonradiatively releasing the energy as heat, or due to FRET with a rate *k_FRET_*. Another source of de-excitation from *S*_1_ is due inter-system crossing to the triplet state, *S*_1_ → *T*_1_, also known as triplet blinking, with a rate *k_Blinking_*. This transition is rare relative to the *S*_1_ → *S*_0_ transitions, and hence is infrequent. However so is the *T*_1_ → *S*_0_ transition (called here de-blinking), with a rate *k_De−blinking_*. In summary, although blinking is rare, when it occurs it takes a long period of time to de-blink and in this time the fluorophore cannot be excited. Although triplet blinking can occur both in the donor and in the acceptor, triplet blinking of the donor does not result in photons. In this analysis we re-color existing photons, therefore processes that do not lead to the emission of a photon will be treated as occurring but not affecting the existing photons we re-color. On the other hand, triplet blinking of the acceptor lead to periods of time in which only the donor can be excited and FRET does not occur, therefore emitting just donor photons with nanotimes exponentially distributed according to *k_D_* as the exponent. Therefore, only the triplet blinking process of the acceptor reproduces another source of photon re-coloring (in this case re-coloring of acceptor photons as donor ones).

In the Jablonski diagram that depicts the photophysical processes in a FRET measurement (Fig. 1, top), the acceptor can be excited via the FRET mechanism. However, at the excitation wavelength intended for donor excitation, acceptor excitation can also occur directly. The fraction of direct acceptor excitation in well-designed smFRET experiments is small but non-negligible. If the acceptor is directly excited, the Jablonski diagram includes just the photophysical processes in the acceptor (Fig. 1, bottom).

**Figure 1:**
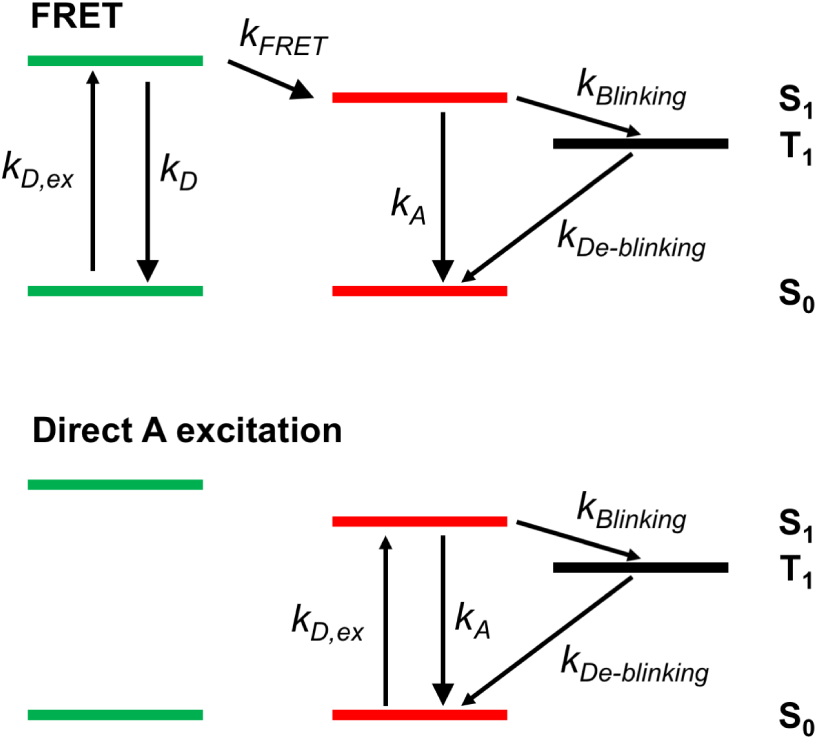
Jablonski diagrams of processes in smFRET leading to photon emission. Top - donor is excited at a rate *k_D,ex_* from ground state *S*_0_ to the first singlet excited state *S*_1_. The donor is de-excited from *S*_1_ back to *S*_0_ either through fluorescence, with a rate *k_D_* (the sum of radiative and nonradiative de-excitation rates), or via FRET, with a rate *k_FRET_* (Eq. 4). The latter leads to an excitation of the acceptor from *S*_0_ to *S*_1_. The acceptor is de-excited either back to *S*_0_, with a rate *k_A_* or to a triplet state, *T*_1_, via inter-system crossing, with a rate *k_Blinking_*. The slow transition from *T*_1_ back to *S*_0_ occurs with a rate *k_De−blinking_*. If the acceptor is in a dark triplet state, it does not function as an acceptor for FRET, causing donor de-excitation only via *k_D_*. Bottom - the acceptor may be excited directly by the excitation source intended for donor excitation.

For a given donor-acceptor distance, *r*, the rate of FRET, *k_FRET_*, depends on the sixth power of *r* and on the Förster radius, *R*_0_ (Eq. 4).

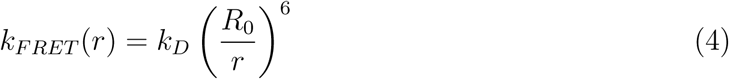

The overall rate of donor de-excitation, *k_D,FRET_*, is the sum of all possible de-excitation processes in the donor (See Fig. 1, top; Eq. 5).

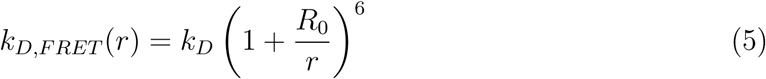

Both *E* and *k_D,FRET_* have probabilistic meanings that are important for the derivation presented in this work. *k_D,FRET_* is related to the probability for donor de-excitation as in Eq. 6.

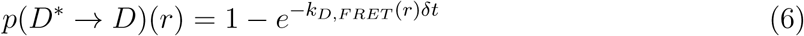

When *r* is constant, Eq. 6 is valid for any *δt* > 0. *E* is the probability that donor de-excitation will occur due to FRET. Therefore, these two parameters are at the heart of simulating the donor excited-state survival dynamics in FRET (Fig. 2). The fundamental experimental observables in smFRET experiments are the identity of the detected photons (D or A photons), the absolute detection times (macrotime), and the detection time relative to the time in which D was excited (nanotime). The distribution of donor or acceptor nanotimes, also known as the donor or acceptor fluorescence decays, describe the characteristic times of the *S*_1_ → *S*_0_ transition. These times can be retrieved from the solution of rate equations resulting from the Jablonski diagram in Fig. 1. Then, the de-excitation rates can be retrieved as the exponent in fits of the fluorescence decays to exponential functions. However, Eqs. 1, 5 describe the process of FRET for a single constant *r*. What if *r* changes as a function of time, *r*(*t*)? Then, each distance, *r*(*t_i_*), introduces different *k_D,FRET_* and *E* values, and hence multi-exponential decays, where each exponent represents the contribution of a given distance value out of many others. Additionally, rapid *r* dynamics introduces an overall decrease in donor nanotimes and increase FRET events.^24,25^ That is because rapid D-A distance fluctuations introduce differences in the D-A distance between the moment of excitation and the moment of de-excitation. While the value of the D-A distance might have been large at the excitation time (hence with an initial low FRET probability), rapid D-A distance fluctuations may lead to a smaller distance value with a higher probability for FRET to occur, hence to a net increase in the apparent FRET efficiency. *r* dynamics can be incorporated into analytical rate equations whose solution can be compared to experiments to find best-fit parameters. Such approach has been used in bulk time-resolved FRET experiments^20,25,26^ but, to the best of our knowledge, never in smFRET. However, this analytical approach becomes impractical as more complex photo-physical schemes are added. Conversely, as shown in the next section, the Monte-Carlo scheme proposed here can be extended to describe more complex photophysics with minimal efforts.

**Figure 2:**
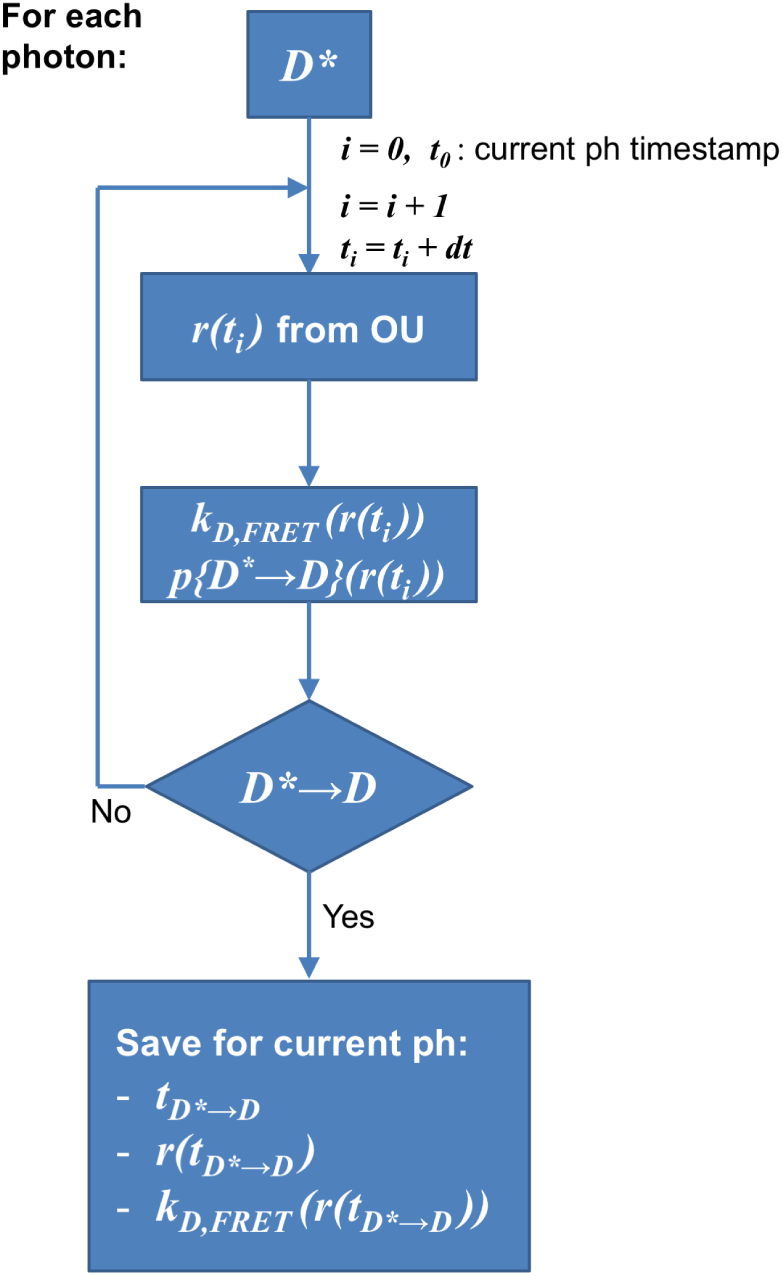
The core algorithm for the Monte-Carlo calculation of FRET per photon. Starting from *t*_0_, immediately after excitation (the donor is excited, *D\**), the donor-acceptor distance, *r*, is advanced in time-steps *dt* (usually of the TCSPC resolution) according to an Ornstein-Uhlenbeck (OU) process of diffusion in a potential well, to produce a distance trajectory, *r*(*t_i_*).^39^ For each time step, the donor de-excitation rate and probability are calculated. Using the donor de-excitation probability the first Monte-Carlo step is to test whether at a given time step de-excitation occurred. If yes, the time iterations halt and the time of donor de-excitation, the donor-acceptor distance and the rate of donor de-excitation at this time are saved and will be used for Monte-Carlo steps in re-coloring this photon.

### 3.1 MC-DEPI photophysics module

As an alternative to using coupled rate equations to describe the Jablonski diagrams of Fig. 1, it is also possible to use a PDA approach using Monte-Carlo simulations to simulate *r* dynamics and the photophysics. This approach depends on the notion that the model that will best describe the experimental results should yield modeled photons with exactly the same features as the experimental ones. In this approach, we simulate photon IDs and nanotimes on top of existing photon macrotimes, namely re-coloring experimental photons. The diagram in Fig. 2 describes an MC-DEPI simulation per a given photon. The core algorithm starts at excitation time *t*_0_, when the donor reaches the excited state *S*_1_ (*D\** being the excited donor). The trajectory of D-A distances, *r*(*t*), is produced by a separate process that we will describe later (see sub-section 3.2 on the dynamics module of MC-DEPI). The probability of donor de-excitation *p*(*D*\* → *D*)(*r*(*t_i_*)) is computed according to Eq. 6 for each distance, *r*(*t_i_*). Then, we test whether de-excitation occurred by comparing *p*(*D*\* → *D*)(*r*(*t_i_*)) to a uniformly random number between 0 and 1. If donor de-excitation did not occur, we advance to the next distance at time *t*_*i*+1_. When donor de-excitation occurs, we save the current time *t_i_* = *t*_*D*\* →*D*_ and the current D-A distance *r*(*t*_*D*\* →*D*_).

Using these three variables we move on in the re-coloring process (Fig. 3). Using Eq. 1 we calculate *E*. *E*(*r*(*t_D* →D_*)) is the probability that donor de-excitation occurred due to FRET, which leads to acceptor excitation, and eventually emission of an acceptor photon (in a simulation of re-coloring existing photons). Therefore, the MC assessment of FRET is a process of re-coloring photons as acceptor photons if FRET occurred, or as donor photons if not. However, *E*(*r*(*t_D* →D_*)) describes the probability for FRET to occur assuming it is the only process that leads to photon re-coloring. In an smFRET experiment there are other processes that dictate photon IDs: (i) direct acceptor excitation (Fig. 1, bottom), in which, with a probability *d_T_*, a photon from the donor laser is absorbed by the acceptor fluorophore instead of from a donor; (ii) the leakage of a fraction *Lk* of donor photons in the acceptor channel; (iii) the *γ* factor bias caused by the different probabilities of detecting donor or acceptor photons, due to different fluorescence quantum yields and detection efficiencies; (iv) acceptor photoblinking (Fig. 1) causing periods of time that will include solely donor photons due to the acceptor being in a dark triplet state. The probability for acceptor blinking, *p_Blinking_*, is constant and depends on the competition between the inter-system crossing *S*_1_ → *T*_1_ and *S*_1_ → *S*_0_ transitions (Eq. 7).

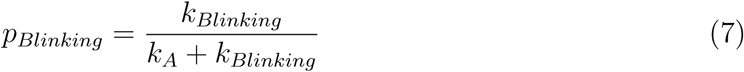

**Figure 3:**
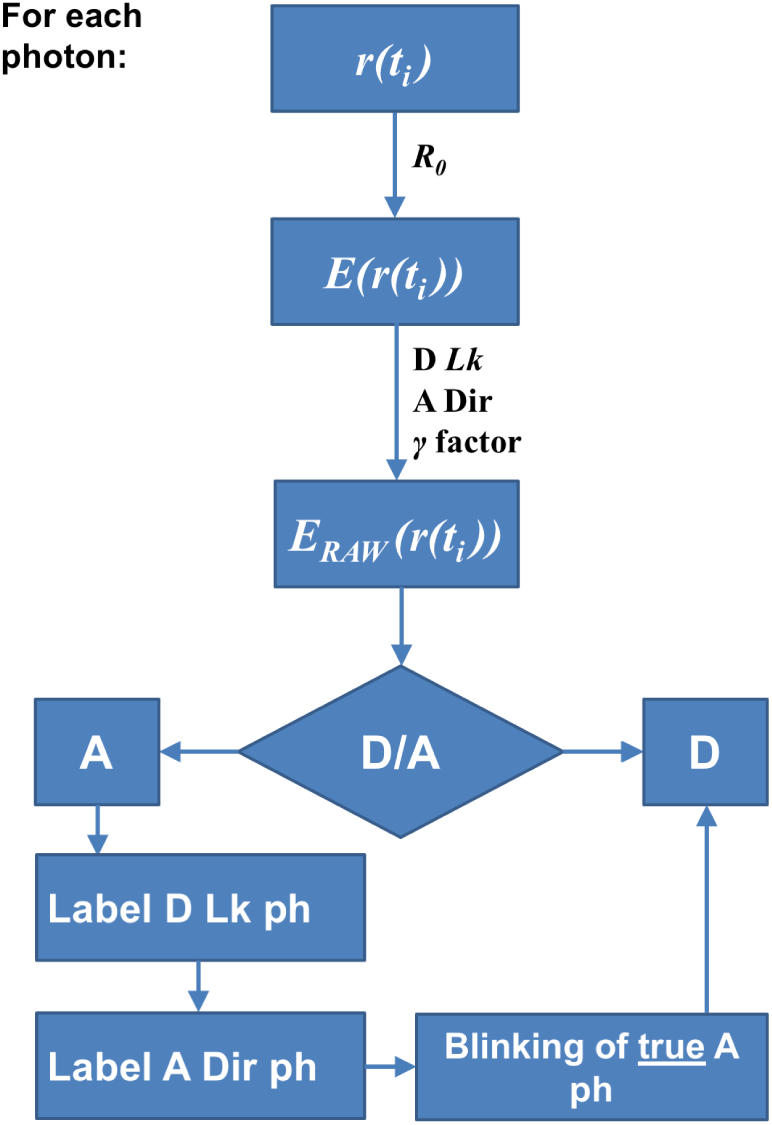
The photon re-coloring scheme per photon. Using the donor-acceptor distance *r*(*t_i_*) at donor de-excitation time saved in the previous step (see Fig. 2), we calculate the FRET efficiency *E*(*r*(*t_i_*)) (Eq. 1). Then, using *γ* factor, the donor leakage *Lk* and the direct acceptor excitation *d_T_* we obtain *E_RAW_* (*r*(*t_i_*)) (Eq. 8). *E_RAW_* (*r*(*t_i_*)) is the probability of detecting a photon in the acceptor channel due FRET, donor leakage, acceptor direct excitation and taking into account the bias introduced by the *γ* factor. Using *E_RAW_* (*r*(*t_i_*)) we randomly label photon as D or A. Next, we randomly select fractions of A photons as leakage and acceptor excitation photons. The remaining A photons are purely due to FRET. We assign A photons from FRET or direct acceptor excitation a nanotime that is the sum of the donor de-excitation time plus a time drawn from the acceptor fluorescence decay distribution. For D and donor-leakage photons, we set the nanotime to the donor de-excitation time. In a last Monte Carlo step, we simulate acceptor photo-blinking. Each *“true”* A photon (due to FRET or direct A excitation) falling during triplet blinking period is relabeled as a D photon. For these photons no FRET can occur, thus the nanotime is drawn from the intrinsic D fluorescence decay distribution. The intrinsic fluorescence decays for both D or A can be single- or multi-exponential.

Normally, in smFRET experiments, the measure *E_RAW_* is corrected to obtain *E*. Here we use the inverse relation (Eq. 8) to compute *E_RAW_* (*r*(*t*_*D*\* →*D*_)) from *E*(*r*(*t*_*D*\* →*D*_)) as a function of the three correction factors *Lk, d_T_* and *γ*. Note that *d_T_* is the ratio of the A and D absorption cross-sections at the donor-excitation wavelength (Eq. 8b). *E_RAW_* (*r*(*t*_*D*\* →*D*_)) is essentially the probability of detecting an A photon with a given FRET efficiency and correction factors. Thus, we use *E_RAW_* (*r*(*t*_*D*\* →*D*_)) to randomly select if a photon is labeled as D or A.

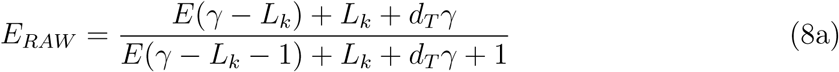

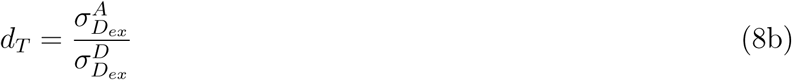

Then, we further label A photons as caused by FRET, direct acceptor excitation or from donor leakage, using the *Lk* and *d_T_*. We remember that only acceptor photons that are labeled as originating from FRET or from direct acceptor excitation are true acceptor-emitted photons. Finally, we simulate periods of acceptor triplet blinking, recoloring acceptor-emitted photons happening during triplet blinking as D photons. The duration of each A dark (blinked) state is drawn randomly from an exponential distribution with mean life-time 1*/k_De−blinking_*. All the acceptor photons re-colored as D due to acceptor triplet blinking, are marked as “acceptor blinked”.

In addition to encoding the photon IDs, we simulate photon nanotimes for different classes of photons as follows: (i) for purely donor photons (not “acceptor blinked”), the nanotime is simply the time of donor de-excitation, *t*_*D*\*→*D*_; (ii) for an acceptor photon due to FRET, the nanotime is the sum of *t*_*D*\* →*D*_ with a random number drawn from the intrinsic A fluorescence decay distribution; (iii) for an acceptor photon from direct acceptor excitation, the nanotime is a random number drawn from the intrinsic A fluorescence decay distribution, since it represents an acceptor photon without the addition of time spent in the donor; (iv) for an acceptor photon due to leakage of donor into the acceptor detection channel, the nanotime is *t*_*D*\* →*D*_, since it represents a donor-emitted photon that was simply detected in the “wrong” detection channel; finally (v) for a donor photon that has been marked as “acceptor blinked”, the nanotime is a random number drawn from the intrinsic D fluorescence decay distribution, since the acceptor is absent, hence the donor in these times acts as a donor-only species. In the simplest case intrinsic D or A fluorescence decays are a exponential distributions with rates *k_D_* or *k_A_*. However, in practice, many organic dyes exhibit more complex decays requiring a multi-exponential model with two or more rates.

### 3.2 MC-DEPI dynamics module

So far we described the photophysics module of MC-DEPI. Now we describe the simulation of trajectories of D-A distances *r*(*t*). Consider a single-molecule burst. The burst is a time series of detected donor and acceptor fluorescence photons. The photon ID, macrotime and nanotime of each photon, described earlier, are accurately defined as:

1. Photon ID of photon *i, ID_i_*, which depends on whether the *i^th^* detected photon in the burst was a donor or acceptor photon
2. Macrotime of photon *i, t_macro_,_i_*, which is the photon detection time at a resolution of a few tens of nanoseconds for the *i^th^* photon in a burst (comparable to pulsed-excitation repetition time in pulsed-excitation smFRET)
3. Nanotime of photon *i, t_nano_,i*, which is the photon detection time relative to the moment of excitation for the *i^th^* photon in a burst (in the resolution of a few ps)

The description of the detection time of the *i^th^* photon in a burst relative to the beginning of the measurement is *t_macro,i_* + *t_nano,i_*. This definition of *t_macro,i_* allows assuming it was the time at which the molecule that produced the detected photon was first excited.

We start by describing *r* dynamics, *r*(*t*), for a single conformational state that is characterized by an equilibrium distribution of donor-acceptor distances, *p_Eq._*(*r*), and distance changes dictated by diffusion in a potential well of the conformational state with a donor-acceptor self-diffusion coefficient, *D*. The relation between the potential well of the simulated conformational state, *U*(*r*), and *p_Eq._*(*r*) is given in Eq. 9.

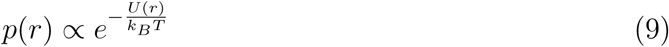

where *k_B_* is the Boltzmann constant and *T* is the absolute temperature. There are many models of *p_Eq._*(*r*), including the Gaussian distribution, *p_G_*(*r*) (Eq. 10a), the skewed or radial Gaussian distribution, *p_rG_*(*r*) (Eq. 10b), and models that describe polymers such as the wormlike chain, *p_WLC_* (*r*) (Eq. 10c).

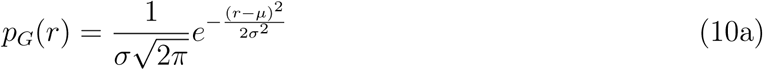

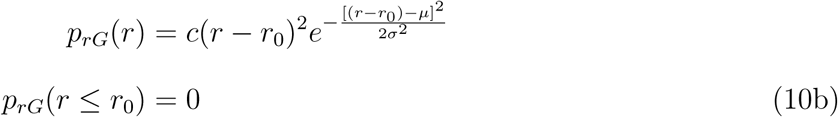

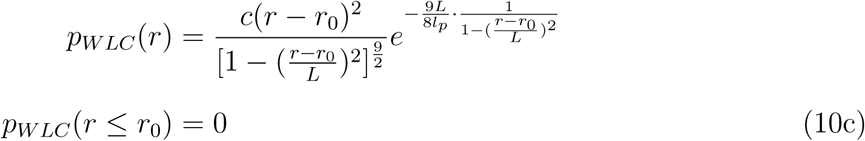

where *c* is a normalization factor, *μ* and *σ* are related to the mean and the standard deviation of the distance (these are exactly the mean and standard deviation of the distance in the case of *p_G_*(*r*)), *L* and *l_p_* are the polymer overall length (the contour length) and the persistence length, respectively, and *r*_0_ is a distance offset below which inter-dye distances do not occur. Models of distance distributions with an offset distance (Eqs. 10b and 10c) are a good approximation for conditions in which dyes are attached to flexible regions of a molecule connected by a rigid part. It is important to note that although *p_G_*(*r*) (Eq. 10a) describes the distribution of distances for a harmonic potential well (see Eq. 9), mathematically it can allow nonzero probabilities for negative distance values, which is of course unphysical. The other distance distribution models described here are mathematically defined to have nonzero probabilities only for distance values above the distance offset, *r*_0_.

While the molecule crossed the detection volume, it had a specific time-trajectory of donor-acceptor distances, *r*(*t*), where at each instance the distance depended on its relevant probability drawn from the distance distribution at equilibrium, *p_Eq._*(*r*), on the distance at the time the previous photon, *i* − 1, was detected and on the time-interval between the previous and the current distances, *t_i_* − *t*_*i*−1_. The dependence *r* in the trajectory, *r*(*t_i_*), on the previous distance, *r*(*t*_*i*−1_), and on the time interval between them changes as a function of the donor-acceptor self-diffusion coefficient, *D* – the faster the diffusional change of *r* is, the faster *r*(*t*) decays from *r*(*t*_*i*−1_) to a distance that is randomly sampled from *p_Eq._*(*r*). For a single-molecule burst, the macrotime of the first photon, *t_macro,_*1, defines time zero for the single-molecule distance trajectory. Therefore the distance at time zero, *r*(*t*_0_), can be drawn randomly from *p*_*Eq*._(*r*). Following a given simulated *r*(*t*), the photon IDs and nanotimes can be simulated from excitation time that yielded the *i^th^* photon to detection time of the *i^th^* photon (from *t_macro,i_* to *t_macro,i_* + *t_nano,i_*) according to the description depicted above (Subsection 3.1 and Figs. 2 and 3). Afterwards, the photon IDs and nanotimes can be simulated from detection time of the *i^th^* photon to excitation time that yielded the *i* + 1^*th*^ photon (*tmacro,i* + *tnano,i* to *t*_*macro,i+1*_). This way, the photon IDs and the nanotimes of the photon time-series are simulated in a photon-by-photon fashion without removing the experimental macrotimes. This is important since the density and number of macrotimes in a burst affect the shot-noise characteristics, the experimental brightness and consequently the time resolution for identification of conformational dynamics. We next move on to describe how to simulate the distance trajectory depending on the underlying *p_Eq._*(*r*) and the self-diffusion coeffcient, *D*.

The following Ornstein-Uhlenbeck (OU) stochastic process describes diffusional motion in a one-dimensional harmonic potential well (Eq. 11).

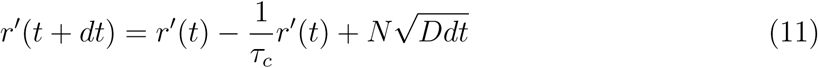

where *N* is a unitary normally distributed random number (mean of 0, standard deviation of 1), *r′*(*t*) is the distance time-series, *dt* is a positive infinitesimal time increment and *τ_c_* is the dynamics relaxation time that depends on the self-diffusion coefficient, *D*, and the standard deviation, *σ*, of the underlying Gaussian distribution, according to Eq. 12.

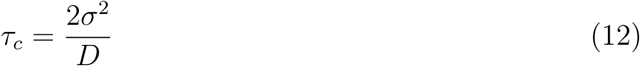

This definition, however, requires using infinitesimally small time increments, *dt*, which makes its use impractical for the typically large time differences between consecutive photons in single-molecule bursts (with intervals in the μs). Gillespie developed a simple direct update formula that allows advancing the OU process in Eqs. 11 and 12 by arbitrary time intervals, Δ*t* (Eq. 13^39^).

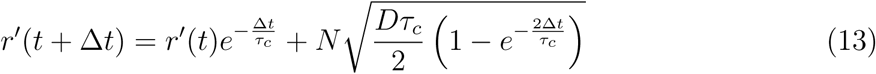

Using this approach allows simulating the distance after a time interval Δ*t*, assuming we know what was the previous distance for a Gaussian distribution with a *μ*=0 and *σ*=1. If *p_G_*(*r*) (Eq. 10a) is the simulated distance distribution and it is defined by other *μ* and *σ* values, each of the simulated distance values, *r′*, can be converted to distance values, *r*, of the underlying distance distribution, which is in this case *p_G_*(*r*), according to Eq. 14.

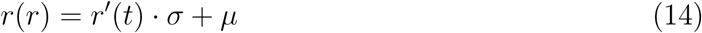

If other models of distance distributions are used, then each distance, *r′*, can be mapped to the distance represented by the simulated distance distribution model, *r*, by considering that the probability of *r′* to occur is the same as the probability of *r* (Eq. 15a).

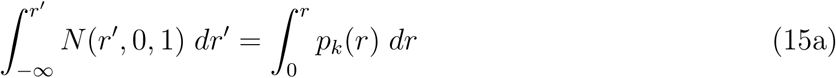

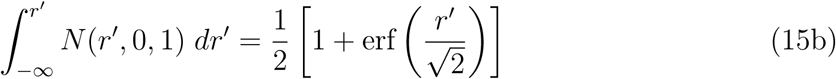

where *p_k_*(*r*) is the simulated distance distribution that might be one of the models presented in Eq. 10 with specific simulated parameter values and *k* represents the name of the model. Note that the integral on the right hand-side of Eq. 15a is taken starting from a distance of 0, since it assumes the model of the simulated distance distribution is defined as a distribution that represents only positive distance values. However the Normal distribution to the left-hand side is symmetric around 0 and defined over the whole real number space. Additionally, an analytical solution to the term on the left (for the standard normal distribution) exists (Eq. 15b). If an analytical solution to the term on the right can be derived, it is preferable to use it explicitly. However if it does not have any known analytical solution, it should be numerically calculated. One can test this procedure to simulate a time-series with constant time steps and show that no matter what values of *p*(*r*) and *D* are used, the autocorrelation of the resulting distance trajectory decays exponentially with a mean relaxation time, *τ_c_*, and that the mean square displacement divided by the time, 〈*r*^2^〉 (*t*)*/t*, as a function of time, *t*, reaches a plateau with a value of the simulated *D*, as expected.

### 3.3 MC-DEPI intra-lifetime diffusion

In time-correlated single photon counting (TCSPC) measurements photon detection times are collected in time bins representing an array of possible discretized nanotimes, also known as TCSPC time bins. The size of the TCSPC bin, *δt*, is typically in picoseconds and defines the accuracy and uncertainty of nanotime recording. The number of TCSPC bins multiplied by the TCSPC bin size (the TCSPC time resolution) defines the maximal possible nanotime values in an experiment. For an experimental array of TCSPC time bins we produce *r′*(*t*) using Gillespie’s direct update formulas for the OU stochastic process (Eq. 13). We start by simulating *r′*(*t* =*δt*) knowing what was *r′*(*t*_0_), which is basically the time of excitation, which we defined as *t_macro,i_*. Then we compute *r′*(*t* = 2*δt*), after a time interval of *δt*, knowing *r′*(*t* =*δt*) that was calculated in the previous step. This stepwise calculation yields the simulated *r′*(*t*) starting from the moment of excitation and throughout all possible TCSPC time bins, in jumps of *δt*. Then, using Eq. 14 or 15 (depending on the underlying simulated *p_Eq._*(*r*)) we map the time-series of distances, *r′*(*t*), that follows a standard normal distribution to a time-series of distances, *r*(*t*), that follows the simulated *p_Eq._*(*r*), using Eq. 15. Next, we follow the steps given above (see subsection 3.1) and in Fig. 2 to calculate the donor de-excitation for the *i^th^* photon, *t*_*D*\*→*D,i*_ and the simulated distance at that time, (*t*_*D*\*→*D,i*_). Then we follow the additional steps given above (see sub-section 3.1) and in Fig. 3 to define the photon ID and nanotime.

### 3.4 MC-DEPI inter-timestamps diffusion

The previous step allowed proper simulation of the photon ID and nanotime of the *i^th^* photon, taking care of possible diffusion-enhanced FRET effects. However, for advancing the simulation to the macrotime of the *i* + 1^*th*^ photon, *t*_*macro,i*+1_, the time of donor de-excitation for the *i^th^* photon, *t_D*→D_,i*, and the simulated distance at that time, *r*(*t_D*_→_D_,i*), is used. The distance that was simulated at *t*_*macro,i*_ + *t_D*→D,i_* and the time interval between this time and the *i* + 1 macrotime, Δ*t* = *t*_*macro,i*+1_ − (*t_macro,i_* + *t_D*→D,i_*) will be used to simulate the distance at *t*_*macro,i*+1_ for the *i* + 1^*th*^ photon using Gillespie’s direct update formulas (Eq. 13) and the distance mapping approach in Eqs. 14 and 15, (both to map distances *r* to *r′* and backwards from *r′* to *r*).

### 3.5 MC-DEPI simulation workflow

The simulation procedure described above will be performed on all bursts, where each burst represents the macromolecule under study that had a different initial distance, *r*(*t*_*macro*,1_), randomly sampled from the underlying simulated equilibrium distance distribution *p_Eq._*(*r*). Due to the burst separation being much larger than the distance fluctuation relaxation time, different bursts have independent initial distance dynamics, representing different molecules out of the ensemble that are not synchronized in time.

The simulation can also describe systems interconverting between more than one conformational state (each associated with a given *p_Eq._*(*r*) and *D*). In this case, the current state is simulated for each photon using a Continuous-Time Markov Chain (CTMC) model. In the CTMC formalism, the probability of being in each state can be computed at arbitrary times knowing the transition matrix and the initial state probabilities. This property is used to randomly generate the current state for each photon. The theory has been described by Gopich and Szabo^41^ with reference to single-molecule experiments or, for the general CTMC theory, in any statistics book treating stochastic processes.^42^ See also the attached notebook “Continous-Time Markov Chain” for details on the formalism. Once the state of the photon is selected, the simulation proceeds as for the single state case, using the distance distribution and diffusion coefficient of the selected state.

### 3.6 MC-DEPI fitting the experiment

Using MC-DEPI, we can compare a simulation of a given set of conditions (*p_Eq._*(*r*), *D*, number of interconverting states and their interconversion rates) to the experimental results. Then, we can iterate the simulation until the simulated results match the experimental ones. The most important observable to be compared is the FRET histogram, similarly to what is done in PDA. But, even when comparing only FRET histograms, MC-DEPI crucially differs from PDA because it takes into account not only distance distributions but also rapid D-A distance changes (the D-A self-diffusion) which generates the diffusion-enhanced FRET effect: rapid D-A changes introduce rapid changes in the probability for FRET to occur, which has a net effect of enhancement of FRET efficiencies and shortening donor de-excitation times. In addition, MC-DEPI includes a detailed description of acceptor triplet blinking. Moreover, differently from PDA, MC-DEPI is a photon-by-photon approach and can reproduce fluorescence decays and other histograms which can yield a more informative comparison with experiments.^7,31,32,43,44^ In this work, we focus on comparing the FRET histograms and the donor and acceptor fluorescence decays. Beechem and Haas have shown that a global analysis of both the donor and acceptor fluorescence decays resulting from a time-resolved FRET measurement, increases the accuracy of the retrieved *p_Eq._*(*r*) and *D* parameters.^25^

In order to find parameters that best fit the experiment, we build a loss function (also called cost function) which is smaller the closer simulation is to the experiment. Due to the Monte Carlo nature of the simulation, the loss function has an intrinsic noise, i.e. multiple evaluation of the same point give different results). For this reason, performing a classical gradient-based optimization which requires a deterministic function is unfeasible. Instead, we use a Bayesian global optimization approach, where at each iteration a new simulation is performed and a new statistical model for the loss function is computed. The statistical model, also known as acquisition function, learns a better approximation of the loss function at each iteration. By minimizing the acquisition function at each iteration, we find a new set of parameters which is chosen as the new point to be evaluated for the loss function. In this work, the acquisition function was computed via Gaussian process regression as implemented in the open source *scikit-optimize* python package.^45^

As noted before, the loss function used in this paper is the sum of two components, one assessing the FRET histograms and the other assessing the fluorescence decays. The E component of the loss function is the mean square error of the simulated and experimental FRET histogram. The fluorescence decay component, conversely, is split in two sub-components one for D and one for A decay. For each decay we use a negative log-likelihood function, similar to what is used for fitting fluorescence decays using the maximum likelihood approach.^46^ In order to combine different losses in a single loss function, we normalize each component by the standard deviation of the Monte Carlo noise. We compute the standard deviation empirically by repeating 100 MC-DEPI simulations with the same set of parameters but with different seeds for the random number generator. Finally, the global loss function is computed as the sum the different components divided by their standard deviation. For details of the derivation of the loss function see section SI-1.3.

The excitation impulse response function (IRF) was taken into account by adding to each simulated nanotime a random number distributed as the experimental IRF distribution. The IRF is different for the D or A channel nanotimes.

### 3.7 Ambiguity in FRET histograms

The most common representation of smFRET results is a histogram of FRET effciency values of all the identified single-molecule bursts, better known as FRET histograms. These histograms show Gaussian-like sub-populations of bursts with common mean FRET effciencies, known as FRET sub-populations. The mean FRET efficiency of a single FRET sub-population is a time-average of FRET dynamics caused by changes of donor-acceptor distances that occurred while the single molecule crossed the detection volume (typically in ms). The distance dynamics that is time-averaged and yields the corresponding FRET sub-population can be described relative to the conformational state characteristics, namely the equilibrium distance distribution, *p_Eq._*(*r*), and the inter-dye self-diffusion coefficient, *D*. However, *p_Eq._*(*r*) can have many different shapes, mean distances and widths, which affect the value of the mean FRET efficiency. Additionally, if distance dynamics occur in times comparable or faster than the donor lifetime, 1*/k_D,FRET_*, the combination of *p_Eq._*(*r*) and *D* influences the value of the mean FRET effciency^24,25^.

Another important experimental parameter is the width of a FRET sub-population. These sub-populations will always have a minimal width caused by the calculation of FRET efficiency from bursts having a limited number of donor and acceptor detected photons and from the effect of background photons, to produce the better known shot-noise-limited FRET sub-population. However, widening beyond the shot-noise limit may occur either due to static heterogeneity (two molecular species with distinct FRET sub-populations that highly overlap) or dynamic heterogeneity (one molecular species that interconverts between multiple conformational states with distinct mean FRET efficiencies). Therefore, widening of a FRET sub-population beyond the shot-noise limit entails additional information.

As a special case, distinct FRET sub-populations may be interpreted as different molecular species having different mean FRET efficiencies. Alternatively, they may be interpreted as an outcome of a single molecular species capable of dynamically transitioning between distinct conformational states characterized by two different mean FRET efficiencies, but only if the timescale of the transitions is larger than the characteristic duration of the single-molecule bursts (which reports on the time it took the molecule to traverse the detection volume). However, if the timescale of transitions between the different conformational states is comparable to the single-molecule burst durations, many bursts will include donor and acceptor photons with different mean FRET efficiencies. This is because while the molecule crosses the detection volume, it interconverts multiple times between different conformational states with different mean FRET efficiencies. The outcome is a FRET histogram that includes a sub-population that *bridges* between the FRET sub-populations of the different conformational states. The faster the transition dynamics is relative to the single-molecule burst durations, the larger the amplitude of the *bridge* sub-population will be on the expense of a decrease in the amplitude of the sub-populations of the original conformational states.^10,47^ Finally, if the dynamics is more than ten times faster than single-molecule burst durations, the outcome will be a single shot-noise limited FRET sub-population with a mean FRET efficiency that equals the equilibrium weighted average of the mean FRET effciencies of the underlying conformational states. Overall, single FRET sub-population may be interpreted in many different ways and not necessarily by a single conformational state with a mean FRET efficiency as that of the FRET sub-population. This point is very important in the debate on how to properly and accurately translate smFRET results into distance information, conformational states and their intricate dynamics.

To stress these points we used the MC-DEPI approach to simulate a multitude of different conditions that may lead to the same shot-noise limited FRET effciency with 〈*E*〉 = 0.4 (arbitrarily chosen). For this set of simulations we chose the following set of parameter values: donor and acceptor fluorescence characterized by a single lifetime component having *τ_D_* = 3.8 ns, *τ_A_* = 4 ns, *R*_0_ = 60 Åand TCSPC bin widths of 10 ps. For the sake of simplicity the simulated fluorescence decays were not convoluted with an IRF. Simulations re-colored the existing photon macrotimes of single-molecule bursts in a measurement of a mixture of two dsDNA molecules doubly labeled with a donor, ATTO 550, and an acceptor, ATTO 647N, with 7 and 17 bp separations (which we name d7 and d17; additional information on the measurement and on the burst analysis is given the Materials and Methods appendix SI-1.2).

The simulated photon IDs and nanotimes are then used to plot the underlying *p_Eq._*(*r*) (Fig. 4, Right-top), the simulated distribution of distances at the time of donor de-excitation, *p*(*r*)@*t_deexcitation_* (Fig. 4, Right-center), the simulated FRET histogram (Fig. 4, Right-bottom) and the simulated donor and acceptor fluorescence decays (Fig. 4, Right-top and bottom, respectively).

**Figure 4:**
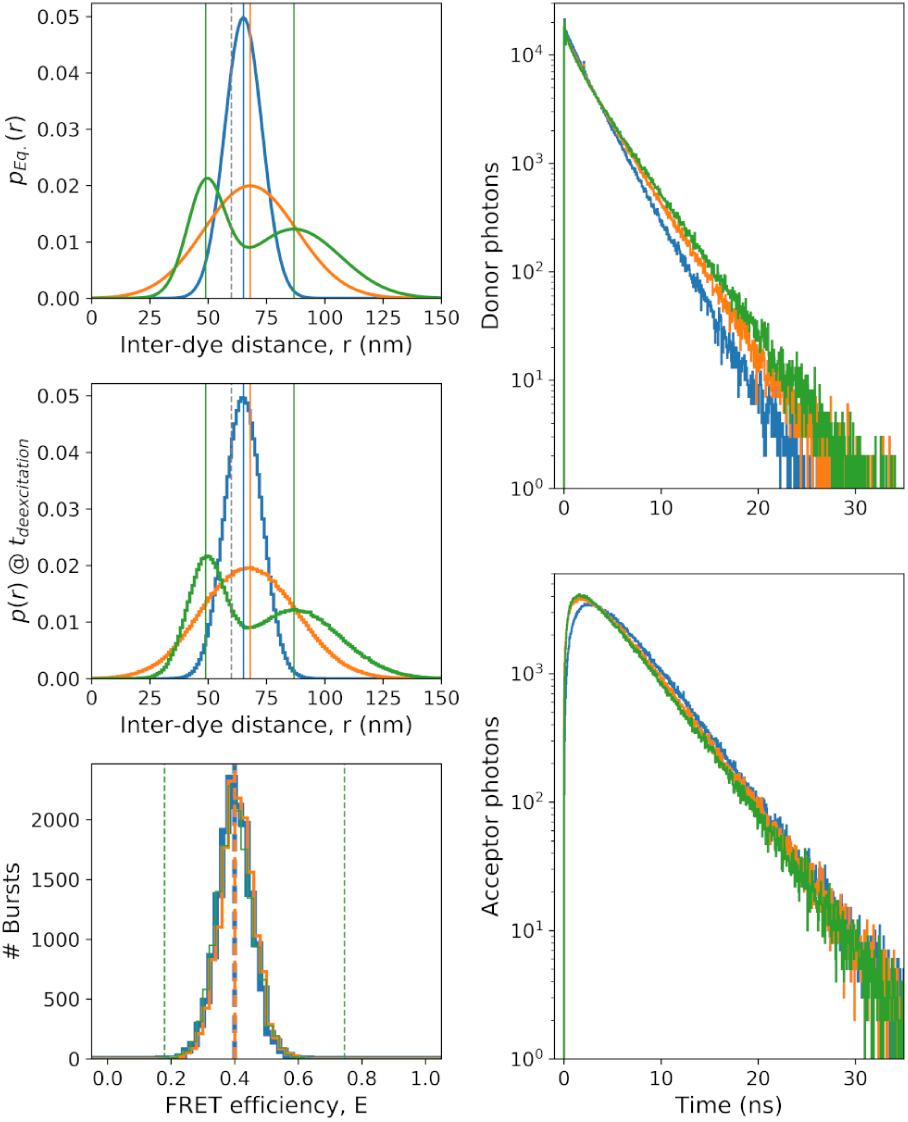
Three different conditions produce the same FRET histograms and the additional information lies in the shape of other histograms. Three different Equilibrium distance distributions, *p_Eq._*(*r*), with different shapes (solid lines) and their corresponding distribution of distances at donor de-excitation time, *p*(*r*)@*t_deexcitation_*, (stair plot). The mean of the underlying *p_Eq._*(*r*) is indicated with vertical lines. This figure reports simulation results of Gaussian *p_Eq._*(*r*) of a single conformational state that is narrow (blue), wide (red) and two conformational states that dynamically interconvert with a relaxation time of *τ_r_*=5 μs (green). A dashed vertical gray line shows the value of *R*_0_. Bottom-left: The FRET histogram of the three different conditions is the same. The dashed vertical lines show the mean FRET values of each state. Right: Donor (top) and acceptor (bottom) fluorescence decays, contain the additional information on the different conditions that yield the same FRET histograms.

As a first example, we simulated three different conditions from conformational states modeled by Gaussian *p_Eq._*(*r*) (Eq. 10a): (i) a single conformational state with a narrow *p_Eq._*(*r*)(*μ*=65.14 Å, *σ*=8 Å, *τ_c_*=50 ns; Fig. 4, blue); (ii) a single state with a wide *p_Eq._*(*r*) (*μ*=68.07 Å, *σ*=20 Å, *τ_c_*=50 ns; 4, orange); and (iii) two conformational states with transition relaxation time of *τ_r_*=5 μs (*f*_1_=0.385, *μ*_1_=49 Å, *σ*_1_=8 Å, *τ_c,_* 1=50 ns, *f*_2_=0.615, *μ*_2_=86.81 Å, *σ*_2_=20 Å, *τ_c_*, 2=50 ns; Fig. 4, green). Note that although these three conditions have very different *p_Eq._*(*r*) (Fig. 4, Left-top), their FRET histograms turn out to be exactly the same, where all characterized by 〈*E*〉 =0.4 and by a shot-noise limited width, even for the case of two conformational states (5 μs transition dynamics results in averaged-out FRET sub-population; Fig. 4, Left-bottom). It is important to note that in the simulated conditions that include FRET dynamics of *τ_c_*=50 ns, the distribution of distances of molecules at the time in which donor was de-excited (Fig. 4, center-left) is the same as *p_Eq._*(*r*) (Fig. 4, top-left; *τ_c_ > τ_D_*, the donor lifetime). However, the donor and acceptor fluorescence decays have different shapes (Fig. 4, Center-top and -bottom, respectively). Therefore for these conditions, the analysis of the fluorescence decays is essential for distinguishing between the three different underlying conditions.

Next we use MC-DEPI to simulate the results of FRET dynamics of a single conformational state modeled by a Gaussian (Eq. 10a), having the same width (*σ*=20 Å) with increasing mean distance (*μ*=68.07, 70.95, 74.5 and 79.28 Å) and decreasing dynamics relaxation time, *τ_c_* (*τ_c_*=50, 8, 2.5 and 0.8 ns; Fig. 5, Left-top, blue, orange, green and red, respectively), where at 8 and 2.5 ns, the dynamics time is comparable to the donor fluorescence lifetime (4 ns) and at 0.8 ns, it is faster than the donor lifetime. In the simulation, the combination of these values correspond to *D* with values of 16, 100, 320 and 1000 Å^2^/ns (using the transformation in Eq. 12). These simulation conditions were chosen to reproduce a FRET histogram with 〈*E*〉 =0.4 (Fig. 5, Left-bottom). Without taking into account the effect of diffusion enhancement on FRET, the increase in distance in these simulations should have yielded a corresponding decrease in 〈*E*〉 (Eq. 1; Fig. 5, Left-bottom, vertical dashed lines). However, if FRET dynamics occurs in times comparable or faster than the donor lifetime (*τ_D_*=4 ns in this case), FRET events from shorter distances represented in *p_Eq._*(*r*) are enhanced due to the rapid dynamics of *r* and the higher *k_D,FRET_* and *p_D_* at lower *r* values (Eqs. 5, 6, respectively). Indeed the distances at the time in which donor de-excitation occurred (Fig. 5, Left-top) are the shorter distances represented in *p_Eq._*(*r*) (Fig. 5, Left-center): the smaller the FRET dynamics relaxation time, *τ_c_*, relative to the donor fluorescence lifetime, *τ_D_*=4 ns, the larger is the deviation between the distribution of distances at the time of donor excitation and at the time of its de-excitation. Therefore, the simulation conditions allow rapid FRET dynamics to compensate for the increase in distances represented in *p_Eq._*(*r*), and to yield the same FRET histograms. If the outcome of these different conditions is the same single FRET sub-population, how can one distinguish between these different conditions? The shapes of the donor fluorescence decays are different (Fig. 5, Right-top). In the case of a wide distance distribution as the one simulated here, if *τ_c_* > *τ_D_*, the donor fluorescence decay will be multi-exponential where each lifetime component represents a different distance out of *p_Eq._*(*r*) (Fig. 5, Right-top, blue). However, from that point on, the smaller *τ_c_* is compared to *τ_D_*, more FRET events occur from shorter distances than from longer ones, which leads to a higher weight of the smaller lifetime components until when *τ_c_ < τ_D_*, the donor fluorescence decay becomes mono-exponential (Fig. 5, Right-top, red). This also affects the acceptor fluorescence decay because it depends on the donor fluorescence decay (Fig. 5, Right-bottom). Therefore, the different shapes of the donor and acceptor fluorescence decays have the additional information that can help in distinguishing between the different conditions simulated here.

**Figure 5:**
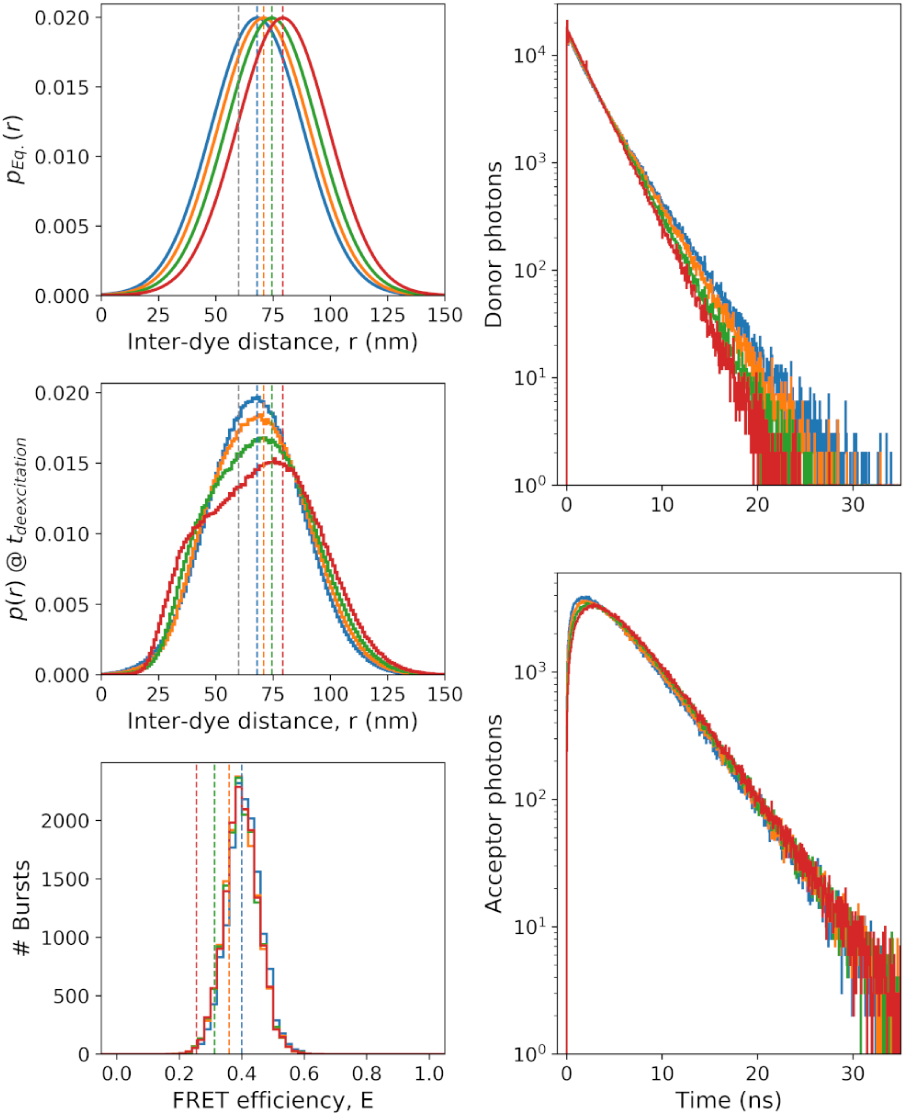
Diffusion-enhanced FRET that lead to the same FRET histogram. Distance dynamics in the timescale (*τ_c_*=50, 8, 2.5 and 0.8 ns as blue, orange, green and red, respectively) of the donor fluorescence lifetime (*τ_D_*=4 ns) lead to the same FRET histogram (Left-bottom) for four different single Gaussian *p_Eq._*(*r*) with the same width and different mean distances (Left-top). *p*(*r*)@*t_deexcitation_* (stair plot) is different than *p_Eq._* (Center-left) due to the effect of enhancement of donor de-excitation at shorter times from shorter distances. The mean of the underlying *p_Eq._*(*r*), without the effect of diffusion-enhanced FRET, is indicated with vertical lines. A dashed vertical gray line shows the value of *R*_0_. The dashed vertical lines show the mean FRET values of each state. Right: Donor (top) and acceptor (bottom) fluorescence decays, contain the additional information on the different conditions that yield the same FRET histograms.

In cases where the sole differences arise from FRET dynamics that take more time compared to *τ_D_*, the fluorescence decays are not expected to include information additional to the FRET histograms (Fig. S1).

Another important set of conditions that may lead to a FRET histogram with a single shot-noise limited FRET sub-population is when there are actually more than a single conformational state and the relaxation time of the transitions between them, *τ_r_*, is slow compared to the donor fluorescence lifetime but much faster than the time the single molecules traverse through the detection volume (Fig. S2).

When the transition dynamics occur in times slower than the single-molecule burst durations (*τ_r_* = 100, 10 ms) the FRET histogram includes two FRET sub-populations that are well-separated. when interconversion dynamics occur in times comparable to burst durations (*τ_r_*=1 ms), a large portion of the bursts include multiple transitions between the two states. Therefore, the FRET **efficiency** values of these bursts are between the values of the mean FRET **efficiency** of the two sub-populations. From that point, the faster the transition dynamics is, the more bursts will include more frequent transitions between the two states, and produce a FRET histogram with a single averaged-out sub-population. Additional experimental information is required to identify that this single FRET sub-population actually represents a time-average of two conformational states with distinct FRET characteristics. The shape of the donor and acceptor fluorescence decays may serve as additional experimental information distinguish between the possibility for a single conformational state and the case of two conformational state, but moving from the former to the latter has to be justified experimentally. One common justification follows statistical inference rules. According to this approach, one moves from a simple model of a single conformational state to a more complex model of two interconverting conformational states only if the former fails to be compared properly to the experimental results.

Finally, acceptor photoblinking can lead to FRET dynamics, between times in which both donor and acceptor photons are being emitted (the FRET species) and others where the acceptor is in a dark triplet state for long periods of time, hence only donor photons are emitted and with nanotimes dictated just by the donor intrinsic de-excitation processes, with a rate *k_D_*. we simulated a set of conditions that may lead to a FRET histogram with a single shot-noise limited FRET sub-population as long as the characteristic times the acceptor spends in the triplet state, *τ_triplet_*, are shorter than the inter-photon times. The simulated conditions were a single Gaussian *p_Eq._*(*r*) (*μ*=65 Å, *σ*=20 Å, *τ_c_*=50 ns) with blinking probability, *p_Blinking_*=0.05 (Eq. 7), and *τ_triplet_*= 5 μs, 250 μs, 1 ms and 5 ms (Fig. 6, Left-top, orange, green, red and magenta, respectively). For comparison, we also simulated another single Gaussian *p_Eq._*(*r*) without acceptor photoblinking, that still lead to a shot-noise limited FRET sub-population with 〈*E*〉 =0.4 (Fig. 6, Left-top, blue). In this simulation the distribution of distances at the time of donor excitation (Fig. 6, Left-top) is identical to the one at the time of donor-de-excitation (Fig. 6, center-left), mainly since the acceptor blinking dynamics is slower than the timescale of the donor lifetime (1*/k_Blinking_, τ_triplet_* >> *τ_D_*). The donor fluorescence decays show slightly different shapes, due to a fraction of the donor photons that originates from a donor-only species when the acceptor is in the triplet state and does not function as an acceptor of FRET (Fig. 6, Right-top). The acceptor fluorescence decays have the same shapes but different amplitudes, since acceptor triplet blinking results in more donor photons on the expense of acceptor photons, since there are long periods in which the acceptor cannot be excited (when it is in the triplet state) and due to the fact that the simulation re-colors a constant amount of photons (Fig. 6, Right-bottom). When acceptor blinking dynamics occur in times slower than the single-molecule burst durations (*τ_blinking_* = 5, 1 ms) the FRET histogram becomes smeared from 〈*E*〉 =0.4 to 〈*E*〉 =0 (Fig. 6, Left-bottom, magenta and red, respectively). when acceptor blinking dynamics occur in times comparable to burst durations but slower than the inter-photon times (*τ_blinking_*=250 μs; Fig. 6, Left-bottom, green), a large portion of the bursts include multiple blinking transitions. Therefore, the FRET sub-population becomes wider than shot-noise limited width. However when *τ_triplet_*=5 μs, the FRET sub-population is characterized by a shot-noise limited width (Fig. 6, Left-bottom, orange). Additional experimental information is required to identify that this single FRET sub-population actually represents a time-average of between a FRET species with 〈*E*〉 > 0.4 and a donor-only species with 〈*E*〉 = 0. The shape of the donor fluorescence decays (Fig. 6, Right-top) may serve as a starting point to distinguish between the possibility for a single conformational state and the case of triplet blinking, but moving from the former to the latter has to be justified experimentally.

**Figure 6:**
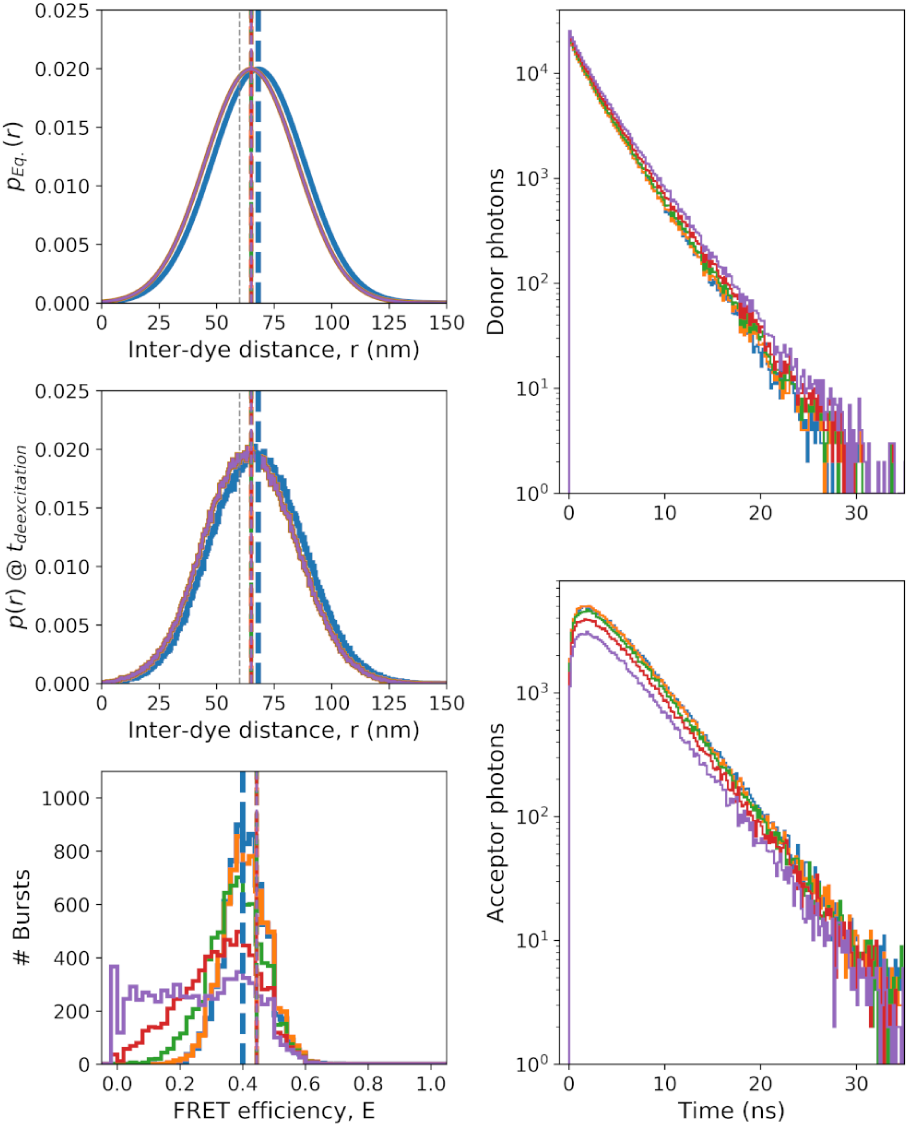
Acceptor blinking dynamics. Acceptor triplet lifetimes in the timescale (*τ_triplet_*=5 μs, 250 μs, 1 ms, 1 ms and 5 ms as blue, orange, green, red and magenta, respectively) with blinking probability, *p_Blinking_*=0.05 (Eq. 7) lead to the same shot-noise limited single population FRET histogram (Left-bottom) for the same single Gaussian *p_Eq._*(*r*) (Left-top, orange, green, red and magenta) as long as *τ_triplet_* is much faster than the inter-photon times (the FRET histogram for *τ_blinking_*=250 μs gets wider and at *τ_blinking_*=1 ms and above becomes smeared from 〈*E*〉 =0.4 dan towards 〈*E*〉 =0) (when the acceptor is blinked, FRET cannot occurs, hence the donor functions as a donor-only species, hence 〈*E*〉 =0 while the acceptor is in the triplet state). *p*(*r*)@*t_deexcitation_* (stair plot) is the same as *p_Eq._*(*r*) (Left-center), because dynamics slower than donor lifetime do not lead to large changes of distances between the times of donor excitation and de-excitation. The mean distance for the underlying *p_Eq._*(*r*) is indicated with vertical lines. A dashed vertical gray line shows the value of *R*_0_. The dashed vertical lines show the mean FRET values of each state. Right: Donor (top) and acceptor (bottom) fluorescence decays of the different simulated conditions have slightly different shapes, hence contain additional information on the different conditions that yield the same FRET histograms. Shown in blue are conditions with which no acceptor triplet blinking yields a shot-noise limited FRET sub-population with 〈*E*〉 =0.4

In summary, the comparison of different experimental histograms (FRET histograms and fluorescence decays) to their MC-DEPI simulated counterparts can serve as a better approach for retrieving the underlying conformational states and their dynamics.

### 3.8 MC-DEPI: the case of double-stranded DNA

Next, we show how by using MC-DEPI simulations we are capable of analyzing smFRET experimental results. For this we performed smFRET measurements of two dsDNA constructs labeled with the same pair of donor and acceptor dyes (ATTO 550 and ATTO 647N, respectively). In one molecule we name d7, the dyes were separated by 7 base-pairs (bp), and in the other we name d17, the dyes were separated by 17 bp. We performed nanosecond alternating laser excitation (nsALEX) smFRET measurements, on freely-diffusing labeled dsDNA molecules, allowing us to: (i) gain the photon ID, its macrotime and its nanotime, for each detected photon; (ii) gain detected photons with interphoton times in the microseconds timescale; and (iii) separate between molecular species with fluorescently active dyes and others where one of the dyes has photobleached. Using a series of control measurements and analyses, we also calculated for each labeled dsDNA molecule the values of *R*_0_, donor fluorescence lifetime components, donor fluorescence quantum yields, and some of the correction factors required for accurate smFRET analysis (for more details on the experiments, please see the Materials and Methods SI-1.2).

First, we analyzed the experimental results of the d7 molecules. We performed a global fit of MC-DEPI simulation results, trying many different conditions, to both the FRET histogram and to the donor and acceptor fluorescence decays. We used the MC-DEPI framework taking into account dsDNA as a single conformational state. In this case, we used a Gaussian distance distribution to describe the conformational state. Additionally, we included the possibility of rapid donor-acceptor self-diffusion to introduce FRET enhancement. We also included acceptor photo-blinking as a possibility. In fitting experimental results to modeled results, the residuals (the difference between the former and the latter) is usually assessed. However, it is important to understand that different simulations of the same parameters induce results that are slightly different from each other. This introduces an intrinsic disper-sion. It is therefore important to understand that the residuals of a given fit are considered satisfactory already if they are comparable to the intrinsic dispersion.

The results of the fitting procedure are shown in Fig. 7. One can see how *τ_c_*=0.9 ns changes the distance distribution from what it is in equilibrium (at excitation time; Fig. 7 Left-top) to what it is at donor de-excitation (Fig. 7, Left-center). The fit to the FRET histogram (Fig. 7, Left-bottom) and to the donor and acceptor fluorescence decays (Fig. 7, Right-top and -bottom, respectively) are shown (Fig. 7, gray - experimental; blue-fitted) following a fit to MC-DEPI simulations with a single Gaussian *p_Eq._*(*r*) with diffusion and acceptor photo-blinking. The best fit parameters of *p_Eq._*(*r*) are *μ*=32 Å and *σ*=15 Å. Using the recently introduced FRET-restrained positioning and screening (FPS) tool,^48^ we calculated the values of *μ* and *σ* expected assuming each orientation in the overall dye available volume is equally probable (Fig. 7, Right-top, inset). The values were *μ*=35.7 Å and *σ*=10.2 Å. Note, however, that the assumptions that this calculation take, may lead to different values. This is because the probability of each orientation in the dye available volume is not necessarily equally probable. Additionally, calculation of the expected FRET using FPS assumes the dye explore all orientations in the dye available volume rapidly relative to the dye fluorescence lifetime. This may not be true in our case. The best fit diffusion relaxation time was *τ_c_*=0.93 ns, which is in the order of the donor fluorescence lifetime (a 0.94 fraction with 4.02 ns and a 0.06 fraction with 0.37 ns). This value together with the value of *σ* translate into a donor-acceptor diffusion coefficient of *D*=484 Å^2^/ns (using the transformation in Eq. 12). A relaxation time of 0.93 ns is well within the dye depolarization times reported for these dyes in the literature.^49–52^ Indeed, FRET is higher than what it should have been without taking into account the diffusion-enhanced effect. However, FRET could have been higher without the balancing effect of acceptor photo-blinking (which reduces FRET values). Without including acceptor photo-blinking, the width of the FRET histogram turns out to be too narrow compared to the experimental one. Additionally, the low FRET tail shown in the experimental results (Fig. 7, Left-bottom, grey) cannot be explained otherwise.

**Figure 7:**
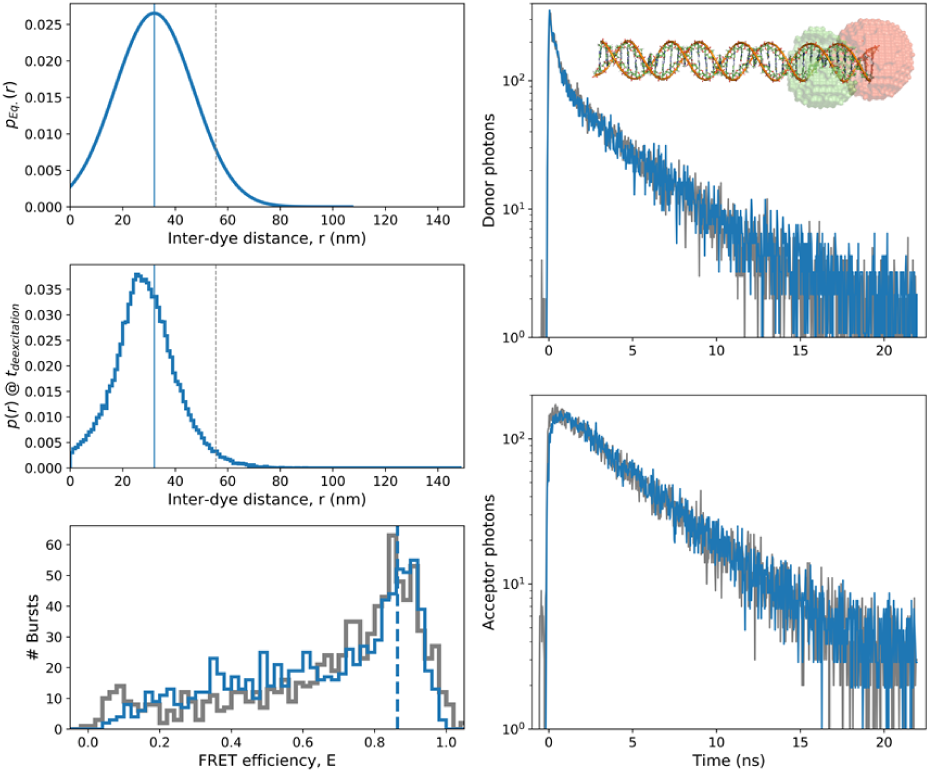
Fitting results to measurements of dsDNA with donor and acceptor separated by 7 bp. nsALEX smFRET measurements of a dsDNA labeled with ATTO 550 and ATTO 647N as donor and acceptor dyes, separated by 7 bp (d7 molecule) were taken. The experimental results are shown in grey and the best fit MC-DEPI simulation of a Gaussian distance distance distribution is shown in blue. The best fit parameters are a single Gaussian *p_Eq._*(*r*) with *μ*=32 Å and *σ*=15 Å, together with distance dynamics *τ_c_*=0.93 ns corresponding to *D*=484 Å^2^/ns (using the transformation in Eq. 12). Additionally, the best fit parameter values of the acceptor photoblinking were *τ_triplet_*=2.04 ms with blinking probability *p_Blinking_*=0.0295. The single Gaussian *p_Eq._*(*r*) is shown (Left-top). Note that the distribution of distances at the time of donor de-excitation, *p*(*r*)@*t_deexcitation_*, is different than the one at the time of excitation (in equilibrium), *p_Eq._*(*r*), mainly at the long distance range, due to the diffusion-enhanced effect. These results fit well with the FRET histogram (Left-bottom) and with the donor and acceptor fluorescence decays (Right-top and bottom, respectively). Note that here the simulated fluorescence decays are shown after convolution with the experimental IRFs. The depiction of the dsDNA molecule with donor and acceptor dyes separated by 7 bp (shown are the dyes available volumes in green and red, respectively), is shown in the inset of the Right-top panel.

The acceptor photo-blinking best-fit parameters are Photo-blinking probability, *p_Blinking_*=0.0295 (which is simply the inter-system crossing efficiency) and an acceptor photo-blinked lifetime, *τ_triplet_*=2.04 ms.

Next we show the best fit results of MC-DEPI simulations for the same dsDNA sample, only this time labeled with donor and acceptor dyes separated by 17 bp (a d17 molecule). This sample includes exactly the same DNA sequence as in the d7 molecule, the same dyes and the same measurement conditions, hence we do not expect to get different acceptor photo-blinking parameters (same photo-blinking probability and same acceptor photo-blinked lifetime). Therefore, we will use the same acceptor photo-blinking parameters found in the fit to the d7 sample, as constants in the fit to the d17 sample.

The results of the fitting procedure are shown in Fig. 8. One can see how *τ_c_*=1.2 ns changes the distance distribution from what it is in equilibrium (at excitation time; Fig. 8 Left-top) to what it is at donor de-excitation (Fig. 8, Left-center). The fit to the FRET histogram (Fig. 8, Left-bottom) and to the donor and acceptor fluorescence decays (Fig. 8, Right-top and -bottom, respectively) are shown (Fig. 8, gray - experimental; blue-fitted) following a fit to MC-DEPI simulations with a single Gaussian *p_Eq._*(*r*) with diffusion and acceptor photo-blinking. The best fit parameters of *p_Eq._*(*r*) are *μ*=71.1 Å and *σ*=15.7 Å. Note that for these *p_Eq._*(*r*) values, if diffusion-enhanced FRET would have been neglected, the mean FRET efficiency would have been lower than what it is with it (Fig. 8, Left-bottom, blue dashed vertical line versus peak FRET population, respectively). Fitting these results without taking into account diffusion-enhanced FRET would have yielded a distance distribution with a significantly shorter mean distance.

**Figure 8:**
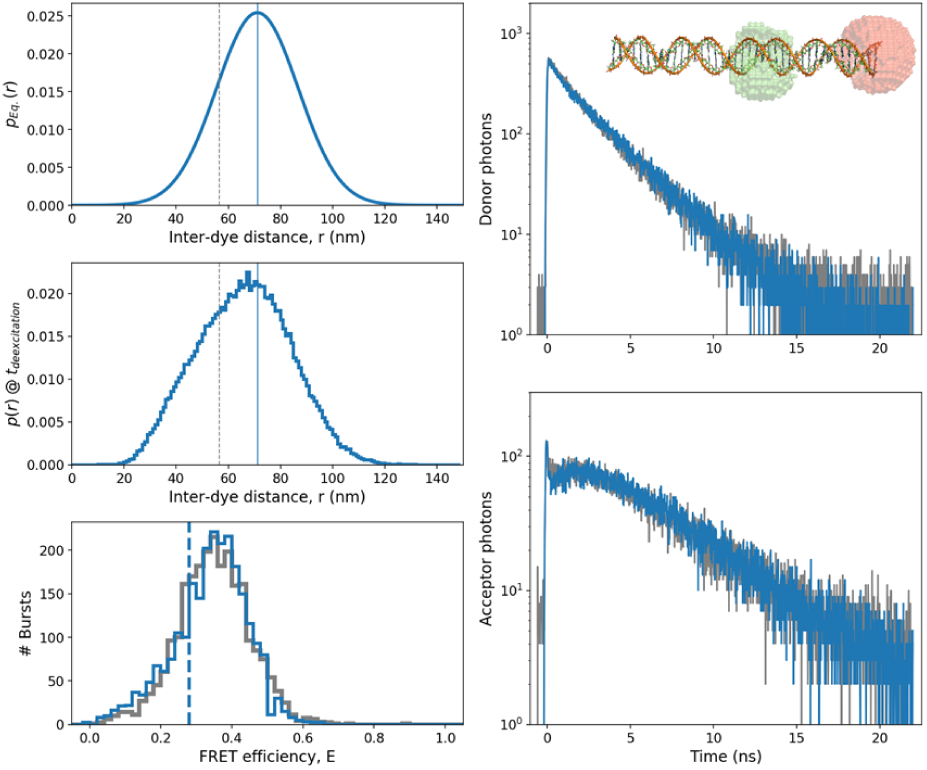
Fitting results to measurements of dsDNA with donor and acceptor separated by 17 bp. nsALEX smFRET measurements of a dsDNA labeled with ATTO 550 and ATTO 647N as donor and acceptor dyes, separated by 17 bp (d17 molecule) were taken. The experimental results are shown in grey and the best fit MC-DEPI simulation of a Gaussian distance distance distribution is shown in blue. The best fit parameters are a single Gaussian *p_Eq._*(*r*) with *μ*=71.1 Å and *σ*=15.7 Å, together with distance dynamics *τ_c_*=1.21 ns corresponding to *D*=407 Å^2^/ns (using the transformation in Eq. 12). The single Gaussian *p_Eq._*(*r*) is shown (Left-top). Note that the distribution of distances at the time of donor de-excitation, *p*(*r*)@*t_deexcitation_*, is different than the one at the time of excitation (in equilibrium), *p_Eq._*(*r*) due to the diffusion-enhanced effect. These results fit well with the FRET histogram (Left-bottom) and with the donor and acceptor fluorescence decays (Right-top and bottom, respectively). Note that here the simulated fluorescence decays are shown after convolution with the experimental IRFs. The depiction of the dsDNA molecule with donor and acceptor dyes separated by 17 bp (shown are the dyes available volumes in green and red, respectively), is shown in the inset of the Right-top panel.

**Figure 9:**
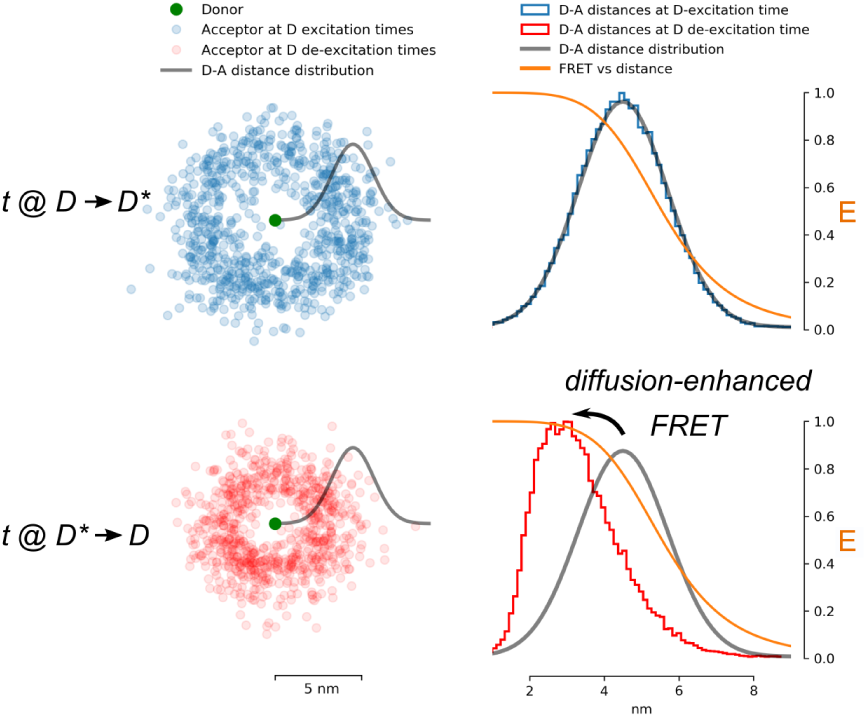
TOC Graphic

Using the recently introduced FRET-restrained positioning and screening (FPS) tool,^48^ we calculated the values of *μ* and *σ* expected assuming each orientation in the overall dye available volume is equally probable (Fig. 8, Right-top, inset). The values were *μ*=65.3 Å and *σ*=10.8 Å. The best fit diffusion relaxation time was *τ_c_*=1.21 ns, which is in the order of the donor fluorescence lifetime (a 0.94 fraction with 3.93 ns and a 0.06 fraction with 0.45 ns). This value together with the value of *σ* translate into a donor-acceptor diffusion coefficient of *D*=407 Å^2^/ns (using the transformation in Eq. 12). A relaxation time of 1.21 ns is well within the dye depolarization times reported for these dyes in the literature.^49–52^ As in the best fit results of d7, also in d17 FRET is higher than what it should have been without taking into account the diffusion-enhanced effect. However, FRET could have been higher without the balancing effect of acceptor photo-blinking (which reduces FRET values). Without including acceptor photo-blinking, the width of the FRET histogram turns out to be too narrow compared to the experimental one.

In summary, using MC-DEPI simulations we were able to fit nsALEX smFRET experimental results of d7 and d17 molecules. Doing so we found that dye linker dynamics, leading to rapid donor-acceptor distance changes (in the range from hundreds of picoseconds to a few nanoseconds) exists and affects FRET results as so did the existence of acceptor photo-blinking. We assume the distance distribution at equilibrium, *p_Eq._*(*r*), that describes dye movements solely due to dye and linker movements, can be described as a Gaussian function. Using this assumption we find that the mean donor-acceptor distances are different than can be calculated using FPS by 10%. Additionally, we find that the standard deviation of the donor-acceptor distance is larger than the one calculated by FPS, by ~5 Å.

## 4 Discussion

### 4.1 Benchmarking on smFRET of dsDNA molecules - not so straightforward

As one can understand, the analysis of smFRET data to yield meaningful and precise distance information is not so straightforward. First, as the simulations have shown, a single FRET population can have many underlying interpretations, all of which are valid as long as not proven otherwise with additional data from the experiment. In the recent years a part of the single-molecule community has put efforts into standardizing smFRET as a tool for accurate retrieval of distance information.^16^ They have done so by measuring the same dsDNA doubly-labeled samples across different laboratories, benchmarking on the rigidity and known structure of dsDNA. The study has shown comparisons of the donor-acceptor apparent distance, calculated directly from the peak FRET efficiency (found by fitting the FRET histogram) using Eq. 1. The authors of this work assumed (i) the dye rotational rate is much faster than the rate of donor de-excitation due to FRET; (ii) the donor-acceptor distance changes (by diffusion) much slower than the donor de-excitation time. Although these assumptions are explicitly expressed, they are not absolutely valid even for molecules such as dsDNA molecules. The assumption that changes in *r* occur much slower than the donor fluorescence lifetime means that each molecule that was excited had a specific value of *r* at the moment of excitation that has not changed until the donor was de-excited, a few hundreds of picoseconds to nanoseconds afterwards. While in many cases this serves as a useful approximation, in other cases this approximation does not hold anymore. For instance, when using smFRET to measure the distance between donor and acceptor fluorophores labeling a rigid dsDNA molecule, this assumption may break. The organic dyes used in smFRET are large and are connected via long linkers. Just the rotational dynamics of the dyes may introduce distance changes, since rotational dynamics are manifested as changes in dye angles, which after multiplication by the dye size yield changes of the positions of the center of the dyes in space. Since the mean size of these dyes from their attachment atoms to dsDNA bases and until the center of the fluorophore is in the range 10-20 Å, such rotational dynamics may introduce large distance changes, that can yield distance distributions with standard deviations in the range 8-10 Å (calculation performed using the FRET-restrained positioning and screening software^48^). The rotational correlation times of such dyes were assessed in several works from analyses of fluorescence anisotropy decays.^49–52^

The typical rotational correlation times of these dyes are in the range 0.3-1 ns, and the typical fluorescence lifetime of these dyes is in the range 1-4 ns. Therefore, in these cases, *r* changes in the timescale of the fluorescence lifetime or slightly faster. In summary, the effect of 0.3-1 ns rotational dynamics of large dyes (with 1-4 ns fluorescence lifetimes) with large linkers on top of a rigid molecule, lead to diffusion-enhanced FRET that has to be taken into account in the distance interpretation of smFRET measurements of molecules as simple as dsDNA (~10% differences in the mean D-A distance compared to when not taking into account the effect of diffusion-enhanced FRET). Additionally, proper description of the underlying distance distribution and acceptor photo-blinking complicate the interpretation of smFRET data on molecules even as simple as dsDNA and have to be properly handled.

### 4.2 Different types of smFRET experiments that can be analyzed by the DEPI approach

All diffusion-based smFRET measurements produce detected photons, where the photon macrotimes and photon IDs are recorded. smFRET measurements based on continuous-wave (cw) excitation do not produce the photon nanotimes. This, however, does not mean the MC-DEPI approach cannot be used to analyze such experimental results, Still it will be very hard if not impossible to distinguish between different combinations of *p*(*r*) and *D* values. Additionally, in cw-based smFRET, the only histograms that are available for comparison with the simulated photons are the FRET histogram. For MC-DEPI analyses of cw-based smFRET measurements, both the donor and acceptor fluorescence lifetimes can serve either as additional free fitting parameters, or can be assumed to have values as the ones reported in the literature.

Additionally, different types of diffusion-based smFRET analyses can help narrow down the search for the parameter value range that yield simulations that fit well with the experimental results. The analysis of fluorescence correlation functions may allow resolving the relaxation times of the conformational changes, *τ_c_*. This can help in better identifying the correlation term from FRET dynamics, to resolve *τ_c_*. A pulsed-interleaved excitation (PIE; also known as nanosecond alternating laser excitation, nsALEX) allows separating single-molecule bursts of molecules carrying just a donor or just an acceptor. This can help identify the exact donor and acceptor fluorescence lifetimes from the same experiment.

Schuler and co-workers have introduced an approach that combines cw excitation with high time resolution using TCSPC to retrieve the FRET-related fluorescence correlation functions that range from seconds to picoseconds. This approach, better known as nsFCS, allowed identifying FRET dynamics at times that are inaccessible in the conventional setups for measuring detecting fluorescence from single molecules.^34^ The analysis according to the MC-DEPI approach can compare simulated FRET histograms and nsFCS correlation curves. In the case of nsFCS, fluorescence decays are not recorded as TCSPC histograms. Nevertheless, the nsFCS correlation functions include the anti-bunching process that can be modeled as analogous to the fluorescence decays, assuming the excitation rate was low.^53^

Finally, the work presented here was based on the distance analysis of FRET assuming the orientational dynamics does not contribute to changes in FRET. Dye rotational dynamics introduce dynamics both in *r* and *κ*^2^. Gopich and Szabo have shown that in case of a constant *r* and rotational correlation times that are five times smaller than donor fluorescence lifetime, rotational dynamics introduce only minute deviations of the FRET efficiency dependence on κ^2^.^27^ The ratio between the dye rotational correlation time and fluorescence lifetime describes experimental values very well for typical organic dyes used in smFRET.^49–52^ Only at ratios larger than 0.5 this deviation becomes significant. Overall, this means we can safely assume the contribution of rotational dynamics to *κ*^2^ dynamics is negligible and dynamics-enhanced FRET is introduced mostly due to distance dynamics. It is important to mention that this assessment was based on the assumption that the probability density function of *κ*^2^ is the one that yields a mean of 2/3, which is introduced by assuming both donor and acceptor experience rotations in all possible *θ* and *ϕ* angles. In most cases not all *θ* and *ϕ* angles are accessible by the dyes, which may or may not introduce a different mean *κ*^2^ value.

A multi-parameter fluorescence detection (MFD) smFRET experiment involves recording single-molecule photons from four different detection channels: donor or acceptor and also parallel and perpendicular polarizations per each donor/acceptor channel.^31^ This allows calculating not only donor and acceptor fluorescence decays but also the associated fluorescence anisotropy decays. Parameters extracted from analysis of fluorescence anisotropy decays have direct links to the dyes’ orientational dynamics and to the boundary conditions of such dynamics in space, usually treated as a cone in which the dye wobbles.^54^ The results of analyses of the rotational dynamics of the dyes from fluorescence anisotropy decays have direct links to, *κ*^2^, through geometrical considerations. If we were to mimic donor and acceptor fluorescence decays through simulating the photon IDs and nanotimes from first principles and distance dynamics, simulating fluorescence anisotropy decays is also possible from first principles and rotational dynamics, which also affect the instantaneous values of *κ*^2^, hence the values of the donor de-excitation rate constants, *k_D,FRET_*, and FRET effciencies, *E*.^55,56^

### 4.3 MC-DEPI: beyond FRET

MC-DEPI allows to simulate, separately, the dynamics and the photophysics, and then to combine them. In the case presented here, the dynamics of a single donor-acceptor distance was simulated and then used in the framework of the FRET photophysics for proper re-coloring of photons. The algorithm for photon re-coloring was laid down as flow charts (Figs. 2 and 3) describing the basic transitions in the Jablonski diagram (Fig. 1) and their associated dependence on the instantaneous donor-acceptor distance at each time step of the dynamics. Therefore the framework of MC-DEPI can allow implementation of other schemes for use in analyses of complex experiments.

As a first example, one can combine FRET with protein-induced fluorescence enhancement (PIFE).^57^ In PIFE a dye that functions as a molecular rotor upon excitation is used. A common dye used in PIFE experiments is Cy3 that exhibits trans-cis isomerization after it is excited mostly in the trans isoform. However, while de-excitation of Cy3 from its trans isoform mostly results in emission of a photon, de-excitation from the cis isoform is almost absolutely nonradiative. Nevertheless, when Cy3 is physically restricted, (e.g. by exclusion of a nearby bound protein), it becomes more fluorescent due to the inhibition of the trans-to-cis transition, leading to increase in its de-excitation from the trans isoform.^22^ Additionally, PIFE can be treated as another molecular ruler. While the FRET ruler reports on donor-acceptor distances in the range 3-9 nm, PIFE may report on distances of Cy3 from the surface of a bound protein in the range 0-3 nm.^22,57^ Therefore, one may simulate the dynamics of the PIFE-measured distance and then employ a series of calculations and Monte Carlo steps to employ the photophysics of PIFE in each step. In reference to FRET, the combination of FRET and PIFE can be employed by simulating the dynamics of the FRET distance and the PIFE distance and at each time step the donor de-excitation is evaluated. The evaluation of the de-excitation event is now dependent not only on the intrinsic fluorescence and on FRET (see Fig. 1) but also on the excited-state trans-cis isomerization rate, where the excited-state cis isoform is treated as tightly-coupled to the cis ground-state.^22^ It is important to remind that dynamics of both types of distances need not be simulated independently. One may think of a more comprehensive model that describes changes in FRET and PIFE distances as correlated.

This way of thinking can be employed also to design an analytical framework to analyze multi-color multi-distance FRET measurements using MC-DEPI. Overall, the logical separation of the dynamics module from the photophysical module allows to use MC-DEPI as a versatile tool for analysis of a variety of different complex experiments with complex photophysical schemes and multiple reaction or conformational coordinates.

## 5 Conclusion

In this work we presented a framework for both simulating and fitting solution-based sm-FRET data based on fundamental FRET photo-physics and on a Monte-Carlo photon recoloring approach. Crucially, we describe the FRET process with D-A distances varying during timescales of fluorescence decays. This is important because rapidly varying D-A distances are a common occurrences when dyes are attached through linkers. The effect on FRET is an apparent increase in FRET efficiency, leading to bias in distance estimations. The theory of FRET with varying D-A distances has been described before in the context of ensemble^25^ and single-molecule data.^27^ However, to the best of our knowledge, this theory has never been applied to quantify smFRET experiments so far.

Our Monte-Carlo diffusion-enhanced photon inference (MC-DEPI) approach takes into account rapid D-A distance fluctuations, dye photoblinking and transitions between multiple states. MC-DEPI allows simulating, as well as analyzing, single-molecule FRET results for the elucidation of D-A distance distributions. It works by recoloring experimentally-detected photons according to an underlying simulation of D-A distance dynamics and its associated photophysics. Using MC-DEPI simulations, we have shown that a single FRET sub-population might be interpreted in many different ways owing to different possible rapid picosecond-nanosecond D-A distance fluctuations, acceptor photoblinking or the possibility of two (or more) conformational states interconverting in timescales faster than typical burst durations. We have shown that in these simulated cases, the shape of the dye fluorescence decays helps in distinguishing between these different possible interpretations. Finally, we have shown how to use MC-DEPI simulations to analyze experimental nanosecond alternating laser excitation (nsALEX) smFRET data by comparing experimental FRET histograms and donor and acceptor fluorescence decays to ones simulated with different underlying dynamical (the D-A equilibrium distance distribution and the D-A distance self-diffusion coeffcient) and photophysical (excited-state transition rate constants) parameters. By doing so, we managed to estimate the D-A distance distribution between dyes attached to dsDNA.

Our results show that organic dyes typically used in smFRET experiments are large enough to introduce D-A distance dynamics in the hundreds of picoseconds timescale stemming from their rotational dynamics. Additionally, after decoupling these diffusion-enhancement FRET effects, the D-A distance distributions show a discrepancy of ~ 10% in the mean D-A distance, relative to the one predicted by existing FRET modeling tools, as well as a D-A distance dispersion higher by ~0.5 nm from the one predicted by FRET modeling tools. In summary, the development and implementation of MC-DEPI provides an important advancement in the growing field of dynamic structural biology based on smFRET measurements.

## 6 Supporting Information

The Supporting Information includes a thorough description of additional MC-DEPI simulations of different conditions that yield the same FRET histograms. It also includes a full description of the Methods used in this work as well as an appendix thoroughly describing the loss function used in this work for the fitting procedure.

## 7 Acknowledgements

We would like to thank Dr. Xavier Michalet for fruitful discussions. This work was supported by the NIH under grant numbers GM069709 (to Shimon Weiss) and GM095904 (to Xavier Michalet); NSF under grant number MCB-1244175 (to Shimon Weiss).

